# Olfactory bulb tracks breathing rhythms and place in freely behaving mice

**DOI:** 10.1101/2024.11.06.622362

**Authors:** Scott C. Sterrett, Teresa M. Findley, Sidney E. Rafilson, Morgan A. Brown, Aldis P Weible, Rebecca Marsden, Takisha Tarvin, Michael Wehr, James M. Murray, Adrienne L. Fairhall, Matthew C. Smear

**Affiliations:** Department of Neurobiology & Biophysics, University of Washington, Seattle, Washington, United States; Institute of Neuroscience, University of Oregon, Eugene, Oregon, United States; Department of Psychology, University of Oregon, Eugene, Oregon, United States; Department of Biology, University of Oregon, Eugene, Oregon, United States; Department of Mathematics, University of Oregon, Eugene, Oregon, United States

## Abstract

Vertebrates sniff to control the odor samples that enter their nose. These samples can not only help identify odorous objects, but also locations and events. However, there is no receptor for place or time. Therefore, to take full advantage of olfactory information, an animal’s brain must contextualize odor-driven activity with information about when, where, and how they sniffed. To better understand contextual information in the olfactory system, we captured the breathing and movements of mice while recording from their olfactory bulb. In stimulus- and task-free experiments, mice structure their breathing into persistent rhythmic states which are synchronous with statelike structure in ongoing neuronal population activity. These population states reflect a strong dependence of individual neuron activity on variation in sniff frequency, which we display using “sniff fields” and quantify using generalized linear models. In addition, many olfactory bulb neurons have “place fields” that display significant dependence of firing on allocentric location, which were comparable with hippocampal neurons recorded under the same conditions. At the population level, a mouse’s location can be decoded from olfactory bulb with similar accuracy to hippocampus. Olfactory bulb place sensitivity cannot be explained by breathing rhythms or scent marks. Taken together, we show that the mouse olfactory bulb tracks breathing rhythms and self-location, which may help unite internal models of self and environment with olfactory information as soon as that information enters the brain.

## Introduction

Animals actively sample their environment and explore space, even in lab experiments without experimenter-controlled stimuli and rewards (Berlyne, 1966; Buzsáki, 2019; Crowcroft, 1973; DeBose & Nevitt, 2008; Land & Tatler, 2009; Osborne et al., 1999; Renner, 1990; Wang & Hayden, 2021). Sampling sensory stimuli provides the raw material for constructing and updating internal models of self and the environment (Behrens et al., 2018; Keller & Mrsic-Flogel, 2018; O’Keefe & Nadel, 1978; Tolman, 1948; Weber et al., 2019; S. C.-H. Yang et al., 2016). In turn, internal models inform perceptual inferences and predict the consequences of actions (Buzsáki, 2019; Churchland et al., 1994; Diamanti et al., 2021; Kleinfeld et al., 2014; Parker et al., 2020; Saleem & Busse, 2023; Webb, 2004). How do sensory samples influence internal models and vice versa?

Sampling behavior imposes structure on odor encounters (Chaput et al., 1992; Crimaldi et al., 2022; Gomez-Marin et al., 2011; Huston et al., 2015; Ravel & Pager, 1990; Schmitt & Ache, 1979; Vanderwolf, 2001; Wachowiak, 2011). In terrestrial vertebrates, breathing provides olfactory sensory neurons with access to odorants, and, even in the absence of odor stimuli, olfactory neurons in the nose and olfactory bulb synchronize their activity to the respiratory cycle (Ackels et al., 2020; Adrian, 1950; Chaput et al., 1992; Grosmaitre et al., 2007; Kay et al., 1996; Macrides & Chorover, 1972; Onoda & Mori, 1980; Vanderwolf, 2000). Animals actively vary their respiratory rhythms depending on the novelty of odor stimuli (Verhagen et al., 2007; Wesson et al., 2008), task context (Frederick et al., 2011; Kepecs et al., 2007), and behavioral goals (Bensafi et al., 2003; Findley et al., 2021; Halpern, 1983; Liao & Kleinfeld, 2023; Welker, 1964). As with other senses (Di Lorenzo, 2021; Fenk et al., 2022; Gibson, 1968; Hayhoe & Ballard, 2005; Kim et al., 2020; Kleinfeld et al., 2006; Michaiel et al., 2020; Rucci & Victor, 2015; Stapleton et al., 2006; Yarbus, 1967), animals move their olfactory organs in order to acquire chemosensory information (Bhattacharyya & Bhalla, 2015; Catania, 2013; Findley et al., 2021; Jones & Urban, 2018; Liu et al., 2020; Youngentob et al., 1987). Movement and location influence the dynamics of stimulus availability to odorant receptors, so animals need to unify odor-driven activity with internal models of how, when, and where they sample the environment (Gire et al., 2016; Nevitt et al., 2008; Vergassola et al., 2007; Wallraff, 2004). Understanding this reciprocal interaction requires studying the olfactory system during active exploration of space (Barwich, 2023; Jacobs, 2012; Jacobs & Schenk, 2003; Poo et al., 2022).

Here, we investigated how exploratory behavior in task-free conditions influences activity in the olfactory bulb, specifically how spiking activity tracks sampling behavior and place. To isolate these factors from potential stimulus- or reward-driven activity, we recorded neuronal activity in the absence of explicit odor cues, task, or reward structure. We find that the breathing rhythms of freely behaving mice are structured on long timescales, persisting in rhythmic states that can last for minutes. Furthermore, the olfactory bulb tracks these breathing rhythms – a statistical model of movement and breathing rhythm can recover stateful structure in the dynamics of neuronal populations. These population dynamics are clearly manifested at the individual neuron level in “sniff fields”, which describe the dependence of neuron firing on latency relative to inhalation and the instantaneous sniff frequency. These sniff fields demonstrate that ongoing activity of olfactory bulb neurons depends on sniff frequency. Moreover, we find that the olfactory bulb tracks place; many individual neurons are significantly modulated by position in space, and the mouse’s location can be decoded from neuronal populations in the bulb with comparable accuracy to neuronal populations in the hippocampus under the same conditions. Importantly, these place-dependent activity patterns do not depend on scent marks or breathing rhythms. Our results show that the olfactory bulb of freely behaving mice contains information about sampling behavior and place, even in the absence of experimenter-controlled odor cues. Thus the integration of odor information into internal models may begin as soon as olfactory information enters the brain.

## Results

### Breathing rhythms are richly structured during spontaneous behavior

We hypothesize that the ongoing activity of the mouse olfactory bulb (OB) encodes information about action and environment in order to contextualize odor-driven input from the nose (Freeman, 1978). This hypothesis predicts that the OB tracks variables such as behavioral state and place, even in the absence of an experimental task. To capture spontaneous behavior and neural dynamics, we implanted mice (n=4) with intranasal thermistors and silicon electrode arrays in the OB, and tracked their movements in a 40 by 15 cm behavioral arena from video under ambient light (see Methods; Fig 1a). We did not impose olfactory stimuli, task structure, or rewards, so that mice experienced only ambient stimuli and generated spontaneous behavior. Most of our recording sessions included a period of head fixation on a stationary platform for comparison with prior experiments (Shusterman et al., 2011), followed by a freely moving period, and then a second head-fixed period, which lasted between 60-90 minutes in total.

**Figure 1:**
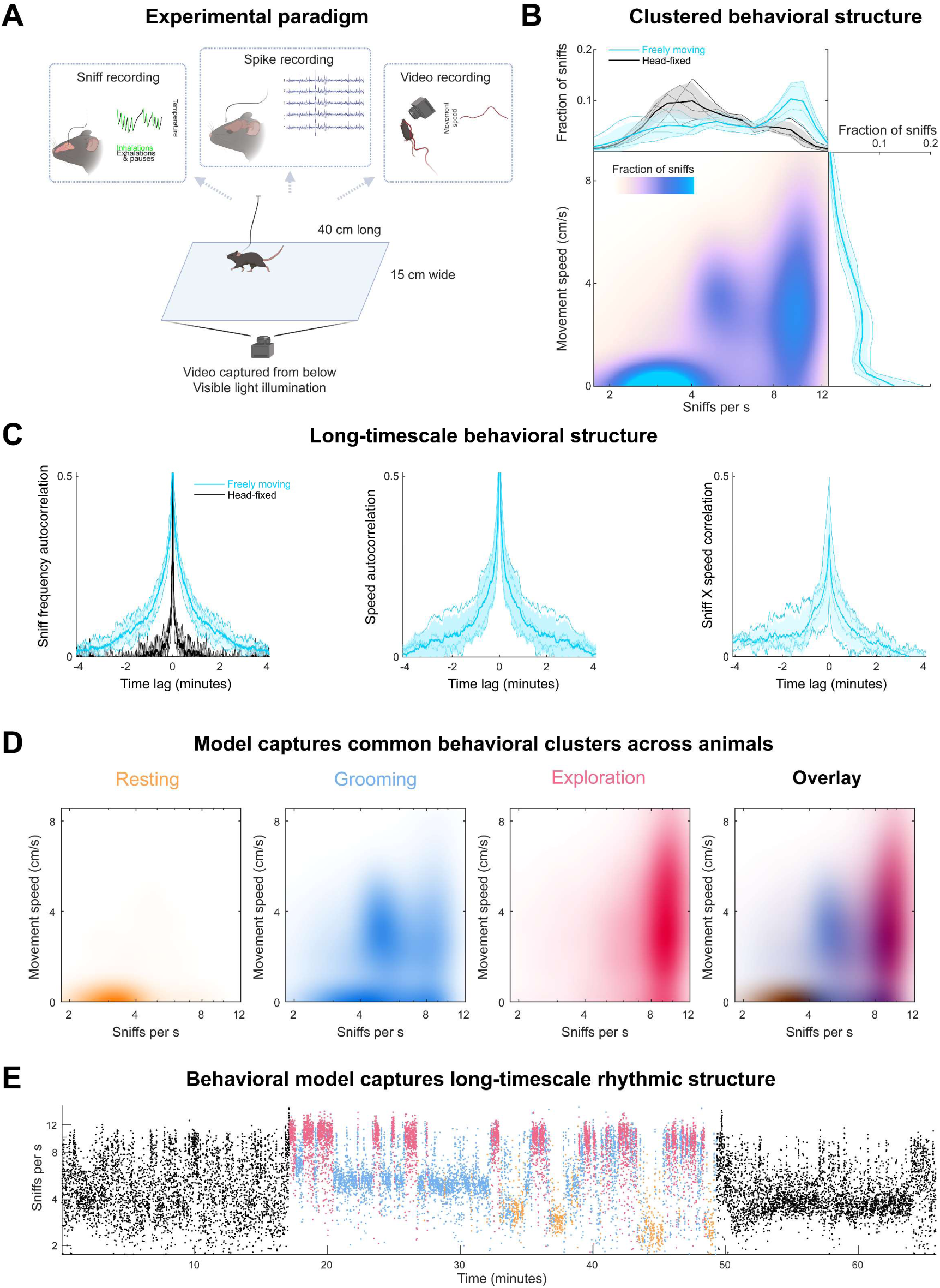
Stateful behavioral structure in an unstructured experimental paradigm. **A.** Experimental setup. Mice were head-fixed or freely moving in a 40 by 15 cm arena while we recorded respiration and neuronal activity and captured video from below in visible light. **B.** The correlation structure of breathing and movement. *Top,* Histogram of instantaneous sniff frequencies of all mice (*n* = 4). Thick lines and shaded regions are mean and ±1 standard deviation, thin lines are individual mice. Blue: freely moving; black: head-fixed. *Right* , Histogram of instantaneous movement speeds, where the movement speed time series was sampled at each inhalation time. *Center,* 2D histogram of breathing frequency and movement speed. **C.** Long-timescale behavioral structure. Autocorrelations of sniff frequency (*Left),* movement speed (*Right)*, and the cross correlation between sniff frequency and speed. Blue: freely moving; black: head-fixed. **D.** A three-state Hidden Markov Model (HMM) fit to the sniff frequency and movement speed time series captures the clustered correlation structure of breathing rhythm and movement. Colormaps show the instantaneous frequency and speed distributions of sniffs in each of three states: Orange: “rest”, blue: “grooming”, red: “exploration”. *Right* Overlay of the distributions from the three states. Overlap is indicated by color mixing and darkness (for colorbars, see Figure 1, supplemental video 2) **E.** The behavioral HMM captures the long-timescale states of breathing rhythms. Each dot indicates an inhalation time with its instantaneous frequency on the vertical axis. Black: head-fixed; other colors as in 1D.

Even in this minimal experimental paradigm, mice exhibited consistently structured behaviors. Mouse breathing is coupled with orofacial and locomotor movements during natural behavior (Findley et al., 2021; Kurnikova et al., 2017; Weinreb, Pearl, et al., 2024). As expected from previous work, breathing rates were overall higher during free movement than during head fixation (Fig 1B, *top*), and breathing rates were correlated with movement speed. In addition to replicating these expected observations, we uncovered novel features of spontaneous behavioral structure. First, we found that instantaneous breathing rates in both conditions were multimodal. During free movement the distribution of sniff frequencies was well fit by a mixture of three log-normal distributions, while during head-fixed conditions, by two (Fig 1B, Fig 1- figure supplement 1). Further, these multiple modes of breathing frequency were associated with distinct movement speeds, such that the joint distribution of sniff frequency and speed formed discrete clusters that recur across sessions and animals (Fig 1B; Fig 1- figure supplement 1). Thus, the relation between sniffing and movement was more complicated than a simple linear correlation.

In addition to the patterning in instantaneous behavior, breathing rhythms and movement speed are structured at longer timescales. The time series of sniff frequency and movement shows stateful organization over timescales of minutes (Fig 1C; Fig 1 - figure supplement 1). In contrast, these persistent states of breathing rhythms are not apparent in head-fixed conditions. To quantify these observations, we computed the autocorrelation of instantaneous breathing frequency and found that the autocorrelation functions had significantly longer timescales in freely moving than head-fixed behavior (Fig 1C). Similar long timescale structure was present in the autocorrelation of movement speed as well as the cross-correlation between breathing and movement. Our analyses demonstrate that even in task-free, ambient-odor conditions, mice perform behaviors structured at multiple timescales.

Continuous, time-varying behaviors can be described as ethograms (Branson et al., 2009; Renner, 2022; Tinbergen, 1965), which divide the time series into discrete behavioral motifs, and provide a useful partition for subsequent analyses (Findley et al., 2021; Markowitz et al., 2023; Weinreb, Pearl, et al., 2024). Motivated by the clustered and long-timescale behavioral structure we observed, we fit the breathing rhythms and movement data from all mice with a Hidden Markov Model (HMM). The model was fit to the behavioral data preprocessed to extract the moving average of speed and the distribution of breathing frequencies, both evaluated in 5 second windows. Model selection was performed using Bayesian Information Criterion (BIC; fig 1 - figure supplement 2). We find that a three-state model well describes free-moving behavioral data and that these three states effectively separate the behavioral clusters (Fig 1D). Through observation of labeled behavioral data, we name these three states “rest”, “grooming”, and “exploration”. These discrete breathing states are seen across all animals but differ in their usage across individual sessions. The average persistence times of states are on the order of tens of seconds to minutes (fig 1 - figure supplement 2). Taken together, these behavioral analyses demonstrate that mice breathe and move in consistently structured ways, even when the experimental paradigm does not impose structure upon their behavior.

### A behavioral model captures the structure of neuronal population dynamics

We next recorded spiking activity in the OB during our task-free, ambient-stimulus paradigm. We recorded from OB with extracellular electrodes, including in our analyses units that passed quality control criteria of fewer than 5 % refractory period violations, and fewer than 10% amplitude cutoff violations (see Methods and Fig 2 – figure supplement 1). Comparing mean firing rates of individual units between head-fixed and freely moving conditions, we found that across the population the mean firing rates were only slightly although significantly different (Fig 2A; head-fixed vs freely moving median = 4.51 vs 5.31 spikes per s; p=0.002, rank sum test ).

**Figure 2:**
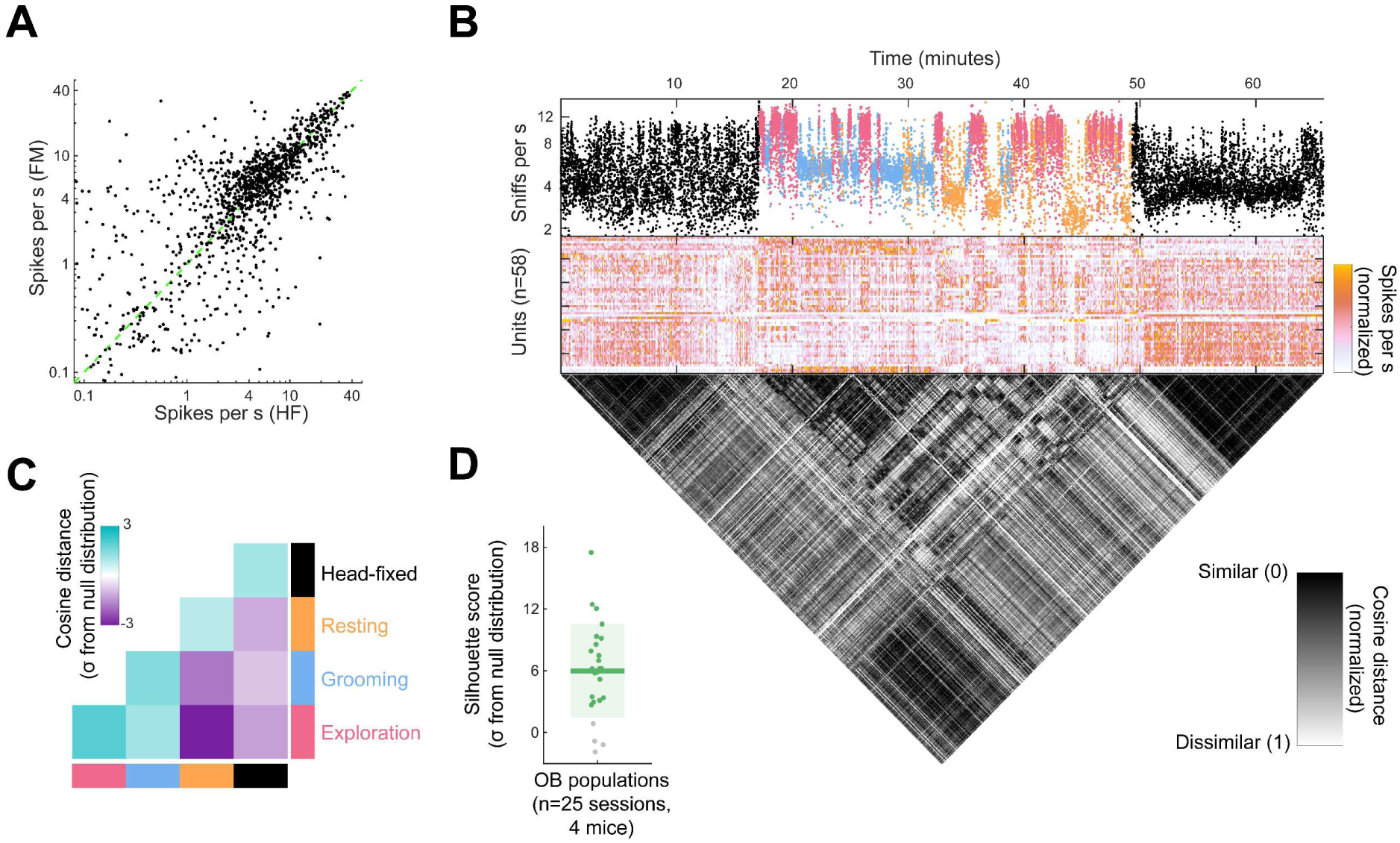
A behavioral model captures the stateful structure of neuronal population activity in Olfactory Bulb. **A.** Scatter plot of mean firing rates during the head-fixed and freely moving epochs of the recording sessions. Each dot indicates the firing rates of an individual unit (n=1680 units in all sessions; n=1274 in sessions with recordable sniff signals). **B.** Behavior, neuronal population activity, and similarity matrix from an individual session. *Top,* Each dot indicates an inhalation time with its instantaneous frequency on the vertical axis. Black: head-fixed; other colors as in 1D. *Middle,* Whole-session spike rates (5 s bins) of the neuronal population recorded in this session. Each row corresponds to an individual unit (n=58 total), with the color scale indicating the normalized firing rates. Each row is normalized separately between minimum and maximum. *Bottom,* Cosine distance matrix quantifies the similarity between the population activity pattern across time bins. **C.** Grand mean cosine distance matrix between states across mice (n=4). Each session’s cosine distance matrix is expressed in units of the number of standard deviations from a null distribution formed by circularly shifting the HMM state time series (see Methods). Positive values indicate greater similarity than expected from the null hypothesis of a “nonsense correlation”; negative indicates less similarity. **D.** Silhouette scores quantifying how well the behavioral states cluster the neuronal population activity patterns in all sessions (4 mice; 25 sessions). Scores are in units of the number of standard deviations from a null distribution formed by circularly shifting the HMM state time series as in 2C.

We predicted that ongoing neuronal activity of OB would reflect the structure of spontaneous behavior. Using the behavioral HMM to partition the sessions, we ask whether behavioral states can describe the similarity of co-occurring population activity. When viewing neural activity alongside behavior, it is apparent that the population vectors are similarly organized into time-varying states (Fig 2B, center). To quantify the similarity of the activity patterns across different time bins in the recording, we computed the cosine distance between population activity in time bins of 5 seconds width, to form a similarity matrix across time throughout a session (Fig 2B, bottom). The apparent block structure of the similarity matrix supports the impression of statefulness in the neural activity. We next compared this structure to the ethograms generated by our behavioral HMM. Importantly, the slow variation in both the behavioral and neural data raises the possibility of a nonsense correlation (Harris, 2021; Meijer, 2021). To quantify similarity relative to that expected from the slow variation in the data, we scored cosine distance as the number of standard deviations away from the mean of a null distribution formed by circularly shifting the time series (see Methods). Taking the grand mean across animals, we find that within a state, activity patterns overlap more than expected under the null distribution, while across states, activity patterns overlap less than predicted by this null hypothesis (Fig 2C). To quantify how well the behavioral HMM clusters the neural data, we calculated a silhouette score, a measure of consistency within clusters, for each session with respect to a circular shift null distribution (see Methods). Most sessions differed significantly from the null prediction (Fig 2D; mean score 6 sigma; 21/25 p<0.01). These analyses show that a model based only on behavioral variables – sniff frequency and movement speed – can effectively cluster neural data from OB, more so than expected from a nonsense correlation arising from the slow variation in behavior and neural activity. Thus, OB activity tracks behavioral structure, even when the behavior is not influenced by experimental stimuli or incentivized by rewards.

### Sniff fields (SnFs) describe how neurons track breathing rhythms

OB activity is already known to be strongly modulated by breathing at the level of individual sniffs (Fukunaga et al., 2012; Macrides & Chorover, 1972; Onoda & Mori, 1980). To visualize the relationship between sniff frequency and unit activity, we aligned spike rasters to inhalation times, and sorted inhalations vertically in descending order of instantaneous sniff frequency (Fig 3A). While most units respond at a consistent latency, some fire with uniform amplitude across sniff frequencies (e.g., Fig 3A, Unit 1), while others fire preferentially during specific frequency ranges, around high (>8 sniff per s; Unit 2), middle (4-8 sniffs per s; Unit 3) or low frequencies (<4 sniffs per s; Unit 4). To capture the joint relationship between inhalation timing and breathing frequency, we describe OB unit activity using “sniff fields” (SnFs), the averaged firing rate as a two-dimensional function of latency from inhalation and instantaneous sniff frequency (Fig 4B). Units displayed a diversity of tuning to frequency, demonstrating that this tuning is not merely a monotonic scaling with sniff frequency. Thus, we observe that variation in breathing rhythm modulates the firing rate of inhalation-synchronized responses in OB units.

**Figure 3:**
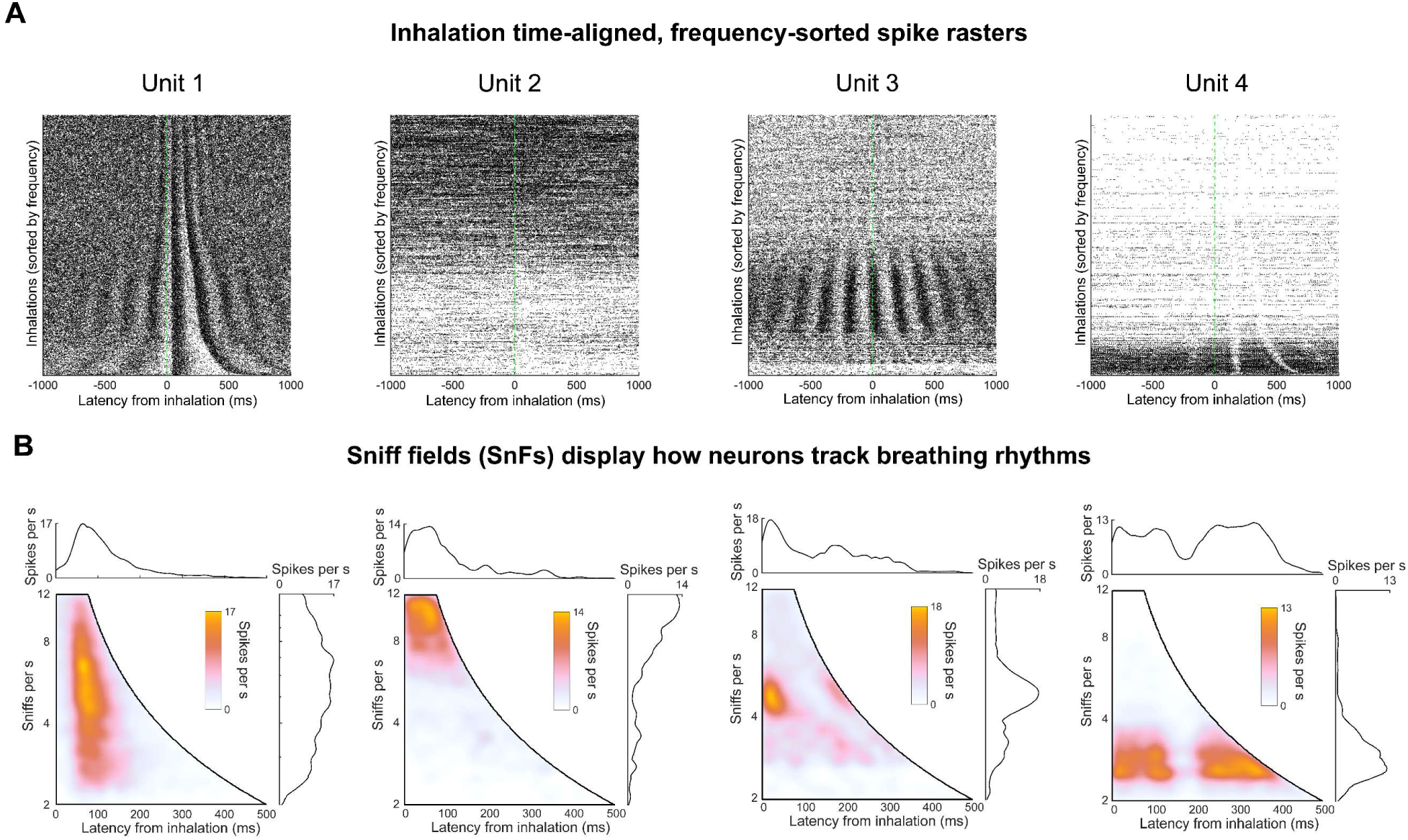
“Sniff fields” (SnFs) display how neurons track breathing rhythms. **A.** Spike rasters from 4 units simultaneously recorded in the same session. Dots indicate spike times relative to inhalation. Each row shows two seconds of the recording centered at each inhalation time at time 0. Rows are sorted in descending order of sniff frequency **B.** Sniff field (SnF) plots from the same four units. *Bottom left,* Colormap indicates firing rates with respect to latency in the sniff cycle and instantaneous sniff frequency. *Right*, Sniff frequency profile of the SnF calculated by taking the max projection across the horizontal axis of the distribution. *Top*, Latency profile of the SnF calculated by taking the max projection across the vertical axis of the distribution.

**Figure 4:**
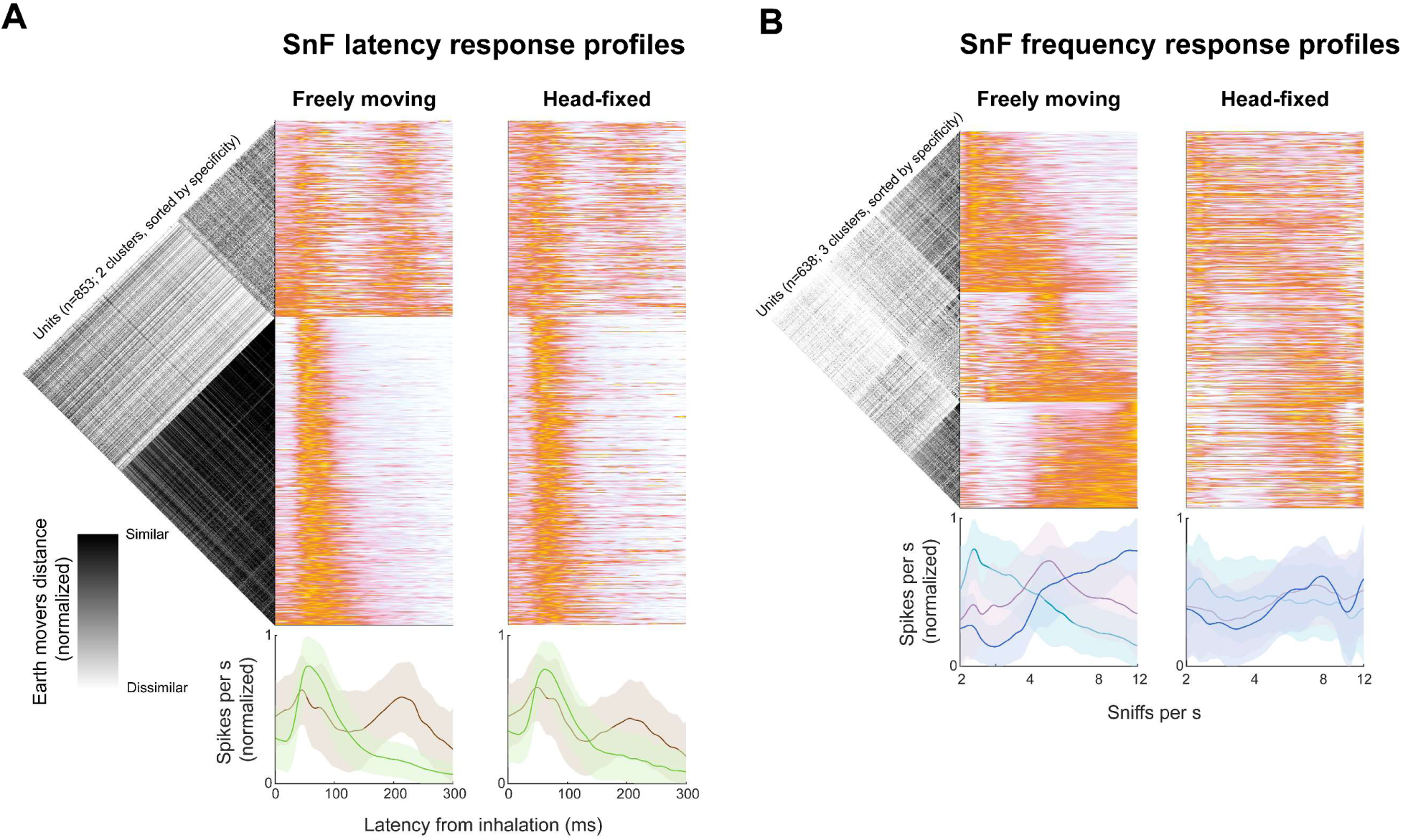
Neuronal sniff field latency and frequency profiles fall into a small number of clusters across the population. **A.** *Right,* SnF latency profiles of all units that were significantly predictable with a GLM fit to spike latency relative to inhalation (n=853/913 p<0.01, Sign rank test). Freely moving and head-fixed matrices are sorted the same. *Left,* units are segregated into two clusters by k-means clustering of the earth mover’s distance matrix quantifying the similarity of SnFs across units. *Bottom,* Within-cluster means for the two clusters. Green: putative tufted cells; Brown: putative mitral cells. Lines and shaded regions are within-cluster means ±1 standard deviation. **B.** *Right,* SnF frequency profiles of all units that were significantly with a GLM trained on instantaneous sniff frequency (n=638/913; p<0.01, Sign rank test). Freely moving and head-fixed matrices are both sorted according to selectivity index in the freely moving data, and differently than the matrices in 4A. *Left,* units are segregated into three types by k-means clustering of the earth mover’s distance matrix quantifying the similarity of SnFs across units. *Bottom,* Within-cluster means for the three clusters. Teal: low frequency units; Purple: medium frequency units; Blue: high frequency units. Lines and shaded regions are within-cluster means ±1 standard deviation.

Across the population, SnFs can be classified into a limited number of types. We extracted SnF latency and frequency response profiles from all units that were significantly predictable by a latency/frequency Generalized Linear Model (GLM) (913/1111 units from sessions with a head-fixed period; p<0.01, sign rank test; see Methods), quantified the similarities of these profiles by calculating earth mover’s distance matrices, and used k-means clustering on these matrices to identify subtypes of SnFs. Among units which significantly encoded latency from inhalation (853/913; p<0.01, Sign rank test), the variety of SnF latency profiles can be captured with two clusters (Fig 4C). One cluster has one peak at <100 ms after inhalation, and another cluster has two peaks, one at <100 ms, and one >200 ms. These two response profiles are consistent with those demonstrated in many studies in anesthetized and awake mammals, and have been shown to correspond with tufted and mitral cell morphology, respectively (Fukunaga et al., 2012; Onoda & Mori, 1980). Separately, among the units that significantly encoded sniff frequency (638/913; p<0.01, sign rank test), clustering the SnF frequency profiles revealed three clusters preferring low, medium, or high sniff frequencies (Fig 4B). The latency and frequency profile subtypes are fairly independent; examples of both latency profile types can be found in all three frequency profile types (Fig 4 – figure supplement 1). Further, instantaneous sniff frequency has a smaller, less consistent relationship with spiking during head fixation than during free-moving conditions (Fig 4B). Taken together, we show that OB neurons not only synchronize their spiking to inhalation, but also track variation in the frequency of the breathing rhythm by varying the amplitude of their inhalation-locked ongoing firing.

### Statistical models reveal that breathing parameters best predict OB activity

To test the extent to which sniff frequency modulation can be explained by inhalation latency modulation, we used a Generalized Linear Model (GLM) to predict individual unit spiking based on behavioral variables. By comparing variables in isolation and in combination, we can ask whether a given variable uniquely contributes to a predictive model of unit firing. We tested models on held out data compared against a null, mean firing rate model by constructing a log-likelihood increase (LLHi) metric (see Methods; (Hardcastle et al., 2017). We perform 10-fold cross validation and calculate statistics on the distribution of LLHi scores. We compared the LLHi scores of GLMs trained on sniff frequency and latency from inhalation, individually and in combination. Consistent with the strong tuning apparent in SnF visualizations, including frequency in the model significantly improves the prediction in 638/913 units, whereas including latency improves the prediction in 853/913 units (Fig 5A). Thus, the OB correlation with sniff frequency is not simply explained by the previously-established synchronization of unit activity to inhalation.

**Figure 5:**
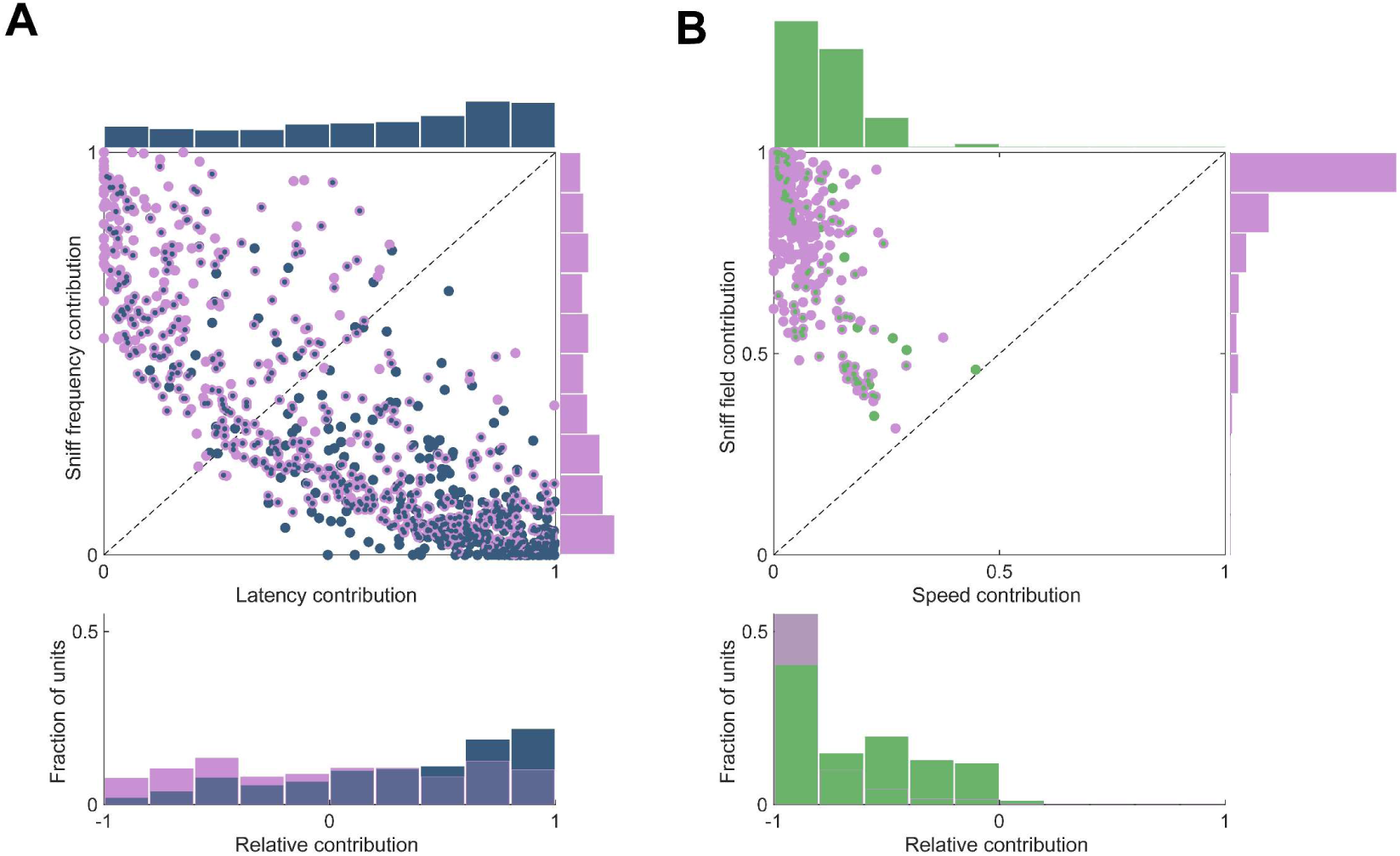
Contribution of behavioral parameters to a predictive model of individual unit firing. **A.** Both frequency and latency improve a model of individual unit firing. *Top,* Each dot indicates a unit with activity that was significantly predictable from a combined GLM based on both latency and frequency (p<0.01, Sign rank test). The contribution is defined as how much including a given parameter improves the model predictions on held-out data (see Methods). Lavender: units for which sniff frequency significantly improved the model prediction; Blue: units for which latency improved the model prediction; Lavender/blue: both parameters improve the model prediction. Marginal distributions of contributions from the two parameters are shown beside and above the scatter plot. *Bottom*, Relative contribution compares the improvement due to the two parameters. **B.** Movement speed minimally improves the predictions of a model incorporating sniff frequency and latency.

We next considered how OB units correlate with movement speed. The behavioral model described above (Fig 1) predicts population activity with both sniff frequency and head movement speed. The correlation between these parameters creates a confounding ambiguity: is the correlation between behavior and neural activity best explained by sniffing, movement, or both? To resolve this ambiguity, we used the same GLM approach. Models based on SnF parameters (a combined frequency/latency model; see Methods) predict firing significantly better than the null model in 913/1153 of units. Models based on movement speed predict unit activity in 249/1153 units (p<0.01, sign rank test), a smaller but still considerable fraction of the population. Thus, as with many other sensory areas of the brain, activity in OB correlates with movement (Parker et al., 2020). However, if we quantify the contribution of these two variables in a combined sniff field/speed model, we find that movement speed improves the predictions relative to that of a model based on SnF parameters in 102/1153 of units (p<0.01, sign rank test; Fig 5B). Further, 12/1153 of the units in our sample were more predictable by speed than by the SnF parameters (Fig 5B *bottom*). Thus, although movement speed is strongly correlated with OB activity, this correlation is largely redundant and reflects more the correlation between breathing and movement than movement itself.

### Olfactory bulb tracks allocentric place

Motivated by the ethological relevance of the relationship between olfactory signals and internal spatial representations (Baker et al., 2018; Gagliardo, 2013; Jackson et al., 2020; Jacobs, 2023; Matheson et al., 2022; Raithel & Gottfried, 2021), and given previous observations of conjunctive odor/place coding in hippocampus (Fischler-Ruiz et al., 2021; Komorowski et al., 2009) and piriform cortex (Kehl et al., 2024; Mena et al., 2023; Poo et al., 2022), we investigated the relation between OB activity and place. Strikingly, individual OB units show apparent place selectivity during free behavior (Fig 6A): spiking activity is spatially modulated for many units. We wondered how this spatial selectivity compared to that of hippocampal neurons, whose place field properties have been extensively studied (Best et al., 2001). However, direct comparison to hippocampal place fields described in the literature is difficult because most recording studies in the hippocampus use experimental paradigms that differ from ours in important ways. First, the behavioral arena we used is smaller than that of most hippocampal studies. Second, hippocampal experiments typically incentivize exploration by distributing food pellets or training on a maze task. Lastly, these experiments often exclude data in which the animals are not moving above a criterion speed. While these design choices successfully establish a focus on canonical place cells, they run counter to the goals of our study. We therefore recorded from neurons in the hippocampus (HPC) of mice in the same arena and task-free experimental paradigm as our OB recordings.

**Figure 6:**
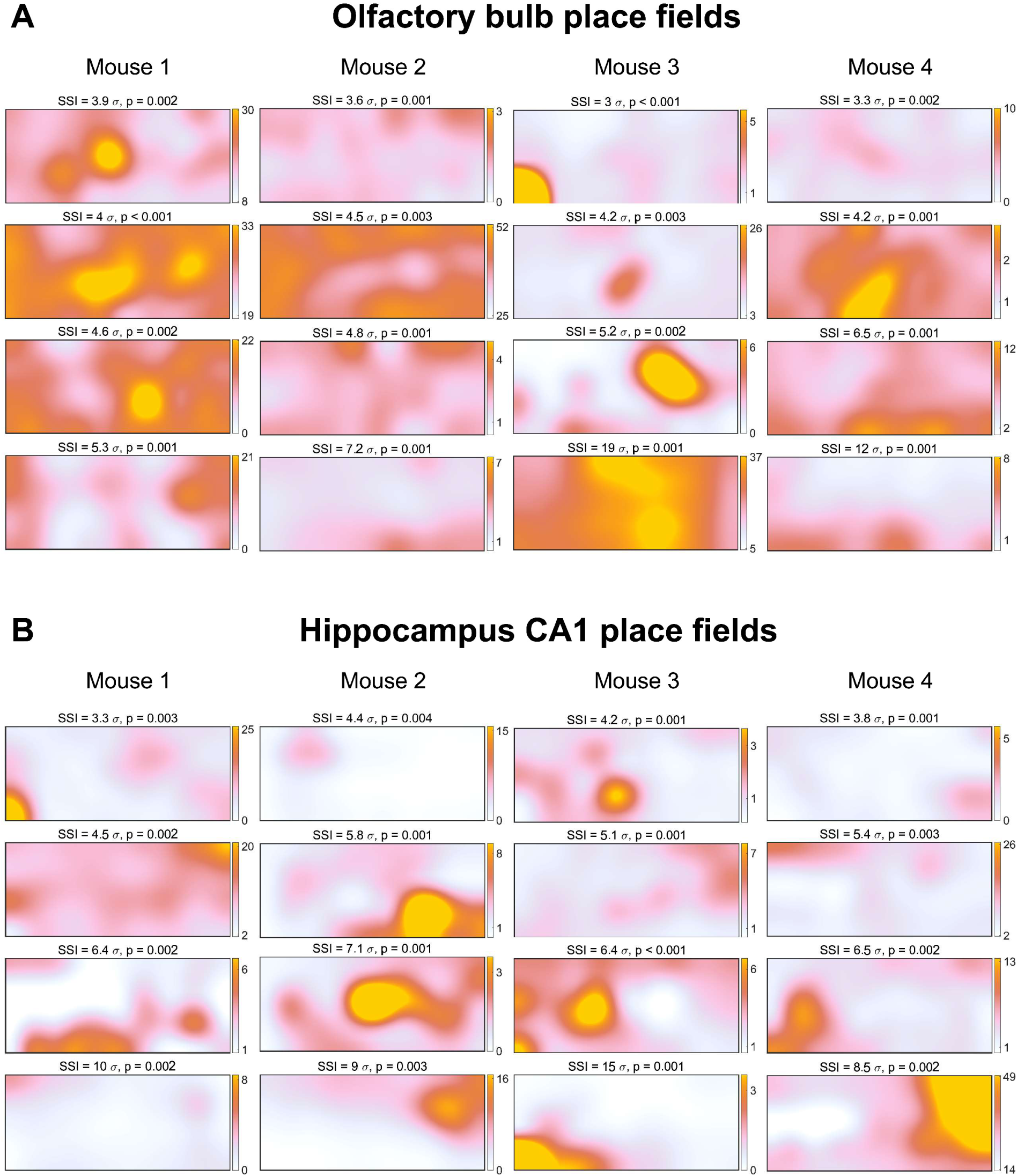
Place fields display allocentric location selectivity of olfactory bulb and hippocampal neurons. Colormaps show occupancy-normalized firing rates as a function of location in the 15 x 40cm experimental arena parsed into a 12 by 5 grid and Gaussian smoothed by one bin width (see Methods). For consistency, OB and HPC colormaps are scaled between the 1^st^ and 99^th^ percentiles of each neuron’s firing rate in 10 s bins. Those values are displayed beside each unit’s colorbar. **A.** Four example OB units from each of four mice. Significance of Spatial Information (SSI) and p-values are defined relative to the circular shift null distributions. **B.** Four example HPC units from each of four mice. OB and HC recordings were performed in different animals.

We compared place selectivity between OB and HPC. Place fields of units recorded from HPC appeared to be more specific than those of OB (Fig 6b, Methods). To quantify the spatial selectivity of individual units in OB and HPC, we modified a traditional metric of place selectivity, spatial information (Skaggs et al., 1992). This information theoretic measure effectively captures the selectivity of canonical place cells, which have very low ongoing firing rates. In contrast, this measure poorly captures the selectivity apparent in neurons with higher ongoing firing rates, such as hippocampal interneurons (Frank et al., 2001; Wilent & Nitz, 2007) or OB neurons (Fig 2A). To better generalize this metric to neurons with ongoing activity, for each unit we compared spatial information to a null distribution formed by circular shifting the position time series 1000 times, and expressed the selectivity as the Significance of Spatial Information (SSI; Stefanini et al., 2020; C. Yang et al., 2024), defined as the number of standard deviations of the real data from the null distribution. This metric, as illustrated by the example cells in Figure 6A, reveals that a substantial minority of OB neurons are spatially selective (196/1557 units, p<0.01), but a significantly smaller fraction than in HPC recorded under the same conditions (270/468 units; Fig 7A).

**Figure 7:**
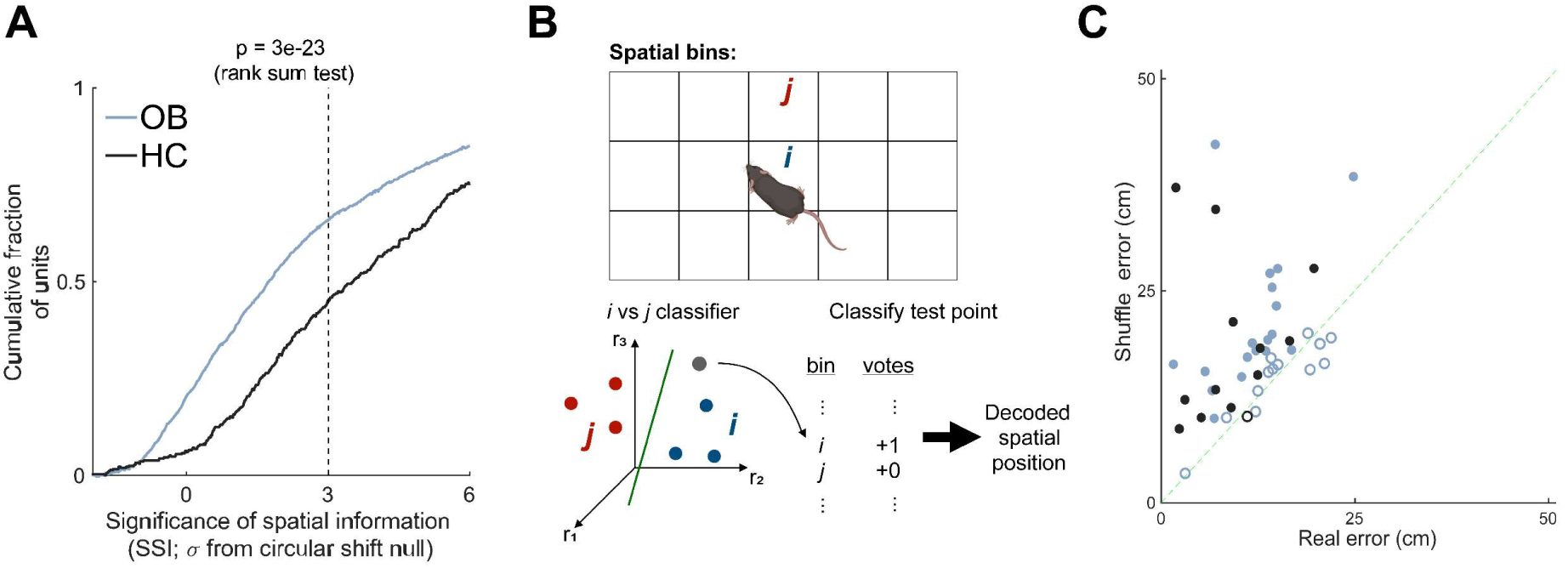
Spatial selectivity of individual neurons and decoding of population activity from olfactory bulb and hippocampus. **A.** Cumulative distributions of selectivity of OB and HPC units, quantified as the significance of spatial information (SSI), defined relative to circular shift null distributions (see Methods). **B.** Decoder model schematic. A classifier for each pair of spatial bins is trained on neuronal activity (firing rate in 5 s bins) and tested on held out data. The decoded spatial position of the mouse at a given time step is taken as the center of the bin that wins the most “votes”, defined as the bin that was predicted by the most pairwise classifiers. **C.** Decoder model performance of OB and HPC populations on real and shuffled controls. Decoding error is defined as the median distance between the decoded spatial position and the mouse’s actual position. Points are individual sessions, filled are p < 0.01 (sign-rank test).

To evaluate spatial information at the level of OB and HPC populations, we trained a decoder model on simultaneous estimates of location extracted from video tracking and population activity, and tested the model’s performance on held-out data at predicting the mouse’s position based on the population activity (Fig 7B; Stefanini et al., 2020). We quantified model performance as the mean error between the decoded position and the actual position (see Methods). For 18/31 sessions from OB and 12/13 sessions from HPC, the model decoded the mouse’s position better than chance (Fig 7C). These analyses demonstrate that in task-free, ambient stimuli conditions, neuronal activity in HPC tracks an animal’s location in an environment. Importantly, we show here, for the first time, that OB neurons also track an animal’s location, consistent with the idea that the olfactory system plays an integral role in navigation (Baker et al., 2018; Dittman & Quinn, 1996; Gagliardo, 2013).

We have shown that OB neurons track breathing rhythms and place. Importantly, breathing rhythms and the states extracted from our behavioral model are not uniformly distributed in allocentric space (Fig 8, figure supplement 1). It is possible that the apparent place information we observe in OB could be explained by differential use of breathing rhythms in different regions in the arena. To test this hypothesis, we used GLMs to ask if inclusion of place significantly improves prediction of single unit spiking activity over a model based on latency from inhalation and sniff frequency (Figs 3 and 4). We found that adding the place covariate to a model with sniff field covariates significantly increased the log-likelihood of held-out data in 160/1153 units (p<0.01, sign rank test), and 81/1153 were better predicted by place than by sniff field (Fig 8A). The unique predictive contribution of place is inconsistent with the hypothesis that place selectivity is an epiphenomenon of the sniff field.

**Figure 8:**
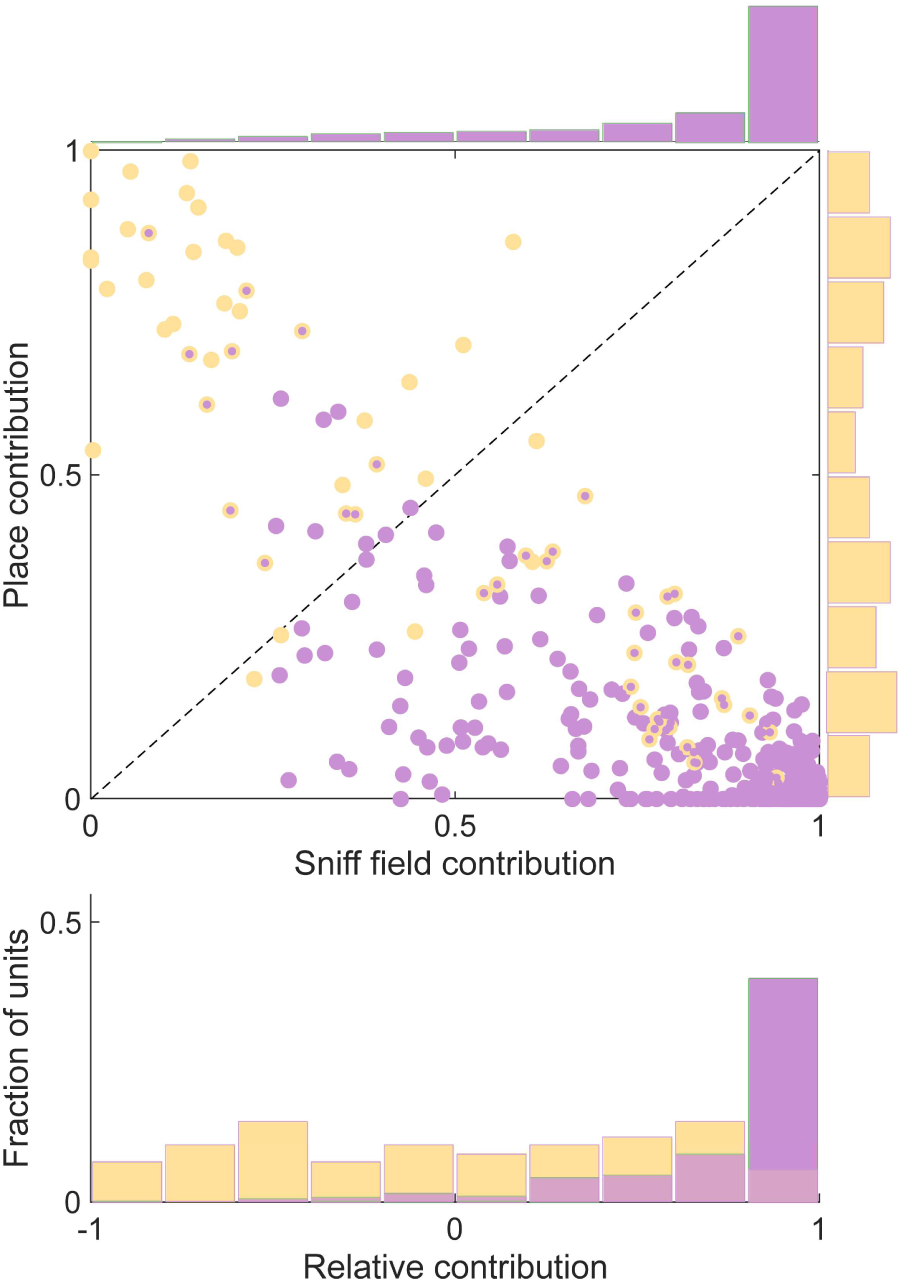
Sniff fields do not explain place selectivity. **A.** *Top,* Each dot indicates a unit that was significantly predictable from a combined GLM based on both sniff fields and place fields (p<0.01, Sign rank test). The contribution is defined as how much including a given parameter improves the model predictions on held-out data (see Methods). Lavender: units for which sniff parameters significantly improved the model prediction; Yellow: units for which place improved the model prediction; Lavender/yellow: both parameters improve the model prediction. Marginal distributions of contributions from the two parameters are shown beside and above the scatter plot. *Bottom*, Relative contribution compares the improvement of the two parameters.

We next considered the possibility that apparent place selectivity could result from responses to ambient odorants. Although we did not apply odor stimuli in these experiments, ambient odors from the mouse and the environment are unavoidable and unevenly distributed in space. The most obvious candidate odor source would be the mouse’s own scent marks: mice, along with many organisms, scent mark their environment, and these marks can contribute to navigational behavior (Drickamer, 2001; Hurst et al., 2001; Khan et al., 2012; Means et al., 1992; Wallace et al., 2002). To test whether scent marks influence spatial selectivity, we performed floor rotation control experiments, in which we rotated a transparent floor mat 180 degrees halfway through a recording session. If place selectivity reflected the location of scent marks, then the “place fields” should rotate to reflect the new distribution of scent marks. Inconsistent with this hypothesis, we observed that place fields often maintained the same location selectivity before and after a floor rotation (Fig 9A). Across the population, the place selectivity of most units maintained a higher correlation across the floor rotation rather than correlating with the new position of the scent marks (Fig 9B,C). Additionally, we asked whether the place decoding models generalize across floor rotation conditions (see Methods). If responses to scent marks drove correlations with place, decoders trained on pre- and tested on post-floor rotation should perform significantly worse than those trained and tested on post-floor rotation data. However, we find that these decoders perform equally well (Fig 9D). Taken together, these findings suggest that place selectivity in the OB does not reflect the distribution of scent marks. However, it is important to recognize that these results do not exclude the possibility that other distal sources of ambient odor explain the place selectivity we observe. In either case, we show that OB contains decodable information about place. Inevitably, this activity will combine with odor-driven activity from the nose and be broadcast to the OB’s numerous postsynaptic targets, most of which send centrifugal feedback to the OB, and several of which are reciprocally connected with the hippocampus (Aqrabawi & Kim, 2018; Padmanabhan et al., 2019; Price, 1985; Reinert & Fukunaga, 2022; Shipley et al., 2008; Vanderwolf, 2001).

**Figure 9:**
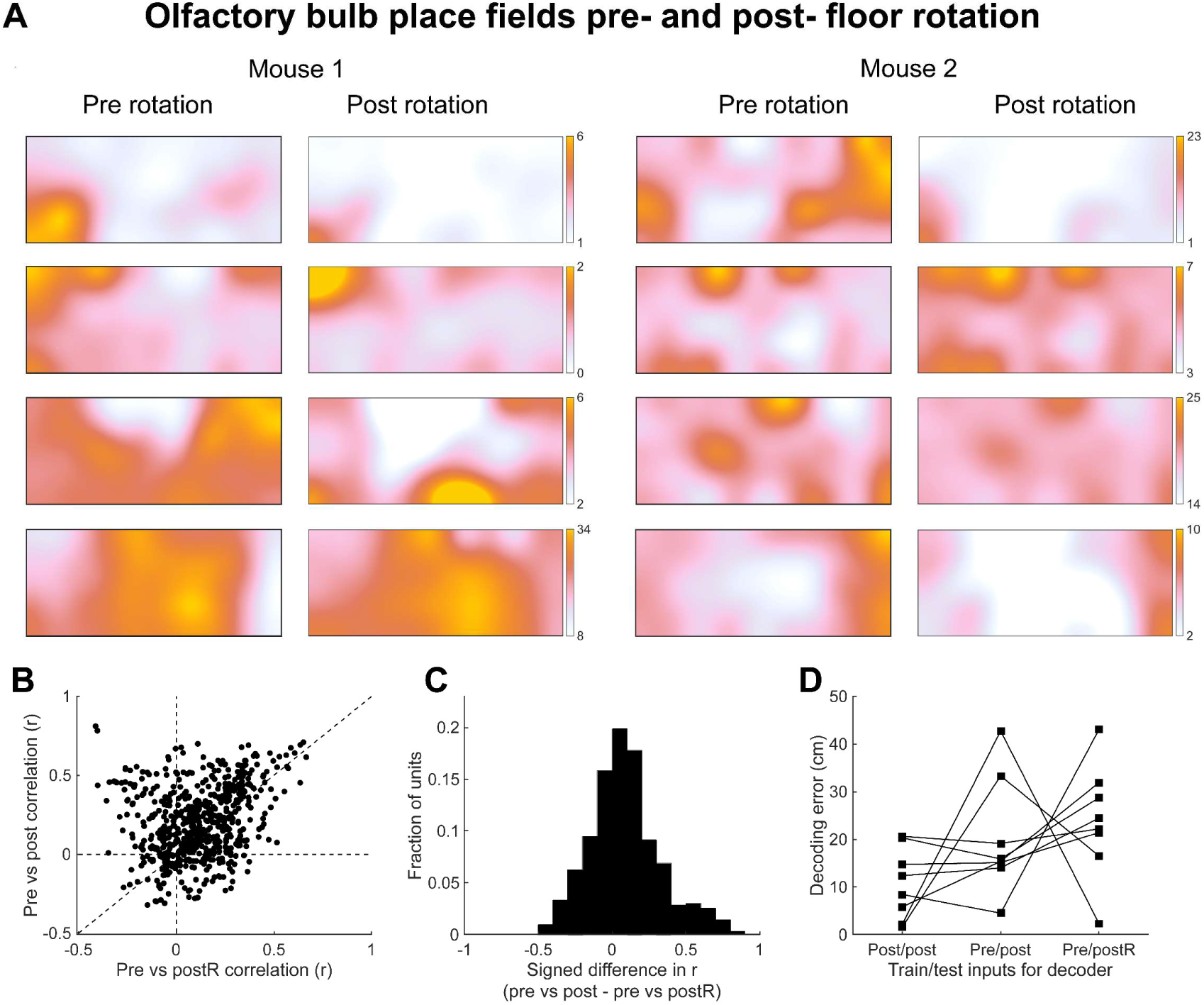
Scent marks do not solely explain place selectivity. In a subset of experiments, we rotated the floor 180 degrees midway through the experiment and compared the resulting place fields. A. Four example units each from OB of two mice. Colormaps are scaled between the 1st and 99th percentiles as in Fig 6. “Pre rotation” shows the place fields calculated from spiking and position time series before the rotation, “post-rotation” shows the place fields from the same units calculated from data after the floor rotation. B. Scatter plot of correlation between pre and post rotation place fields vs pre and virtually rotated post rotation (postR). C. Signed difference in correlations in B reveals that a majority of units’ place fields do not follow the scent marks while others do. D. Population decoding models trained on activity during the post rotation data and tested on post rotation data (Post/post), trained on pre rotation and tested on post rotation (Pre/post), and trained on pre rotation and tested on a 180 degree rotated control of post rotation data (Pre/postR).

## Discussion

In this study, we demonstrate that spontaneous breathing rhythms and olfactory bulb activity share a rich temporal structure in mice. Even during spontaneous behavior and ambient stimuli, mice organize their breathing rhythms into persistent states that evolve in time. Olfactory bulb dynamics are modulated by multiple features of breathing. As previously reported, we show that olfactory bulb spikes synchronize precisely with inhalation onset times, and that this synchronization is very similar between the head-fixed and freely moving states. We also show, for the first time, that many olfactory bulb neurons fire at different rates during different breathing frequencies, and that these activity patterns co-evolve with persistent rhythmic states of breathing behavior.

Further, we find that activity is modulated by the animals’ allocentric location at the single unit and population level. We used the rich data available in free behavior preparations to fit statistical models that tease apart the contributions from these variables and show that the OB multiplexes multiple aspects of the animal’s context. This activity will combine with odor-driven activity from the nose and be broadcast to the OB’s numerous postsynaptic targets, many of which are reciprocally connected with the hippocampus (Aqrabawi & Kim, 2018; Padmanabhan et al., 2019).

The presence of neural correlates, however striking, does not prove that they are adaptively beneficial for the animal (Gould et al., 1979), but the metabolic costs of ongoing activity encourage systems to make use of these representations (Sterling & Laughlin, 2015). Animals sample the same stimuli or environment in different contexts and internal dynamics can help reconfigure representations to be relevant to current demands (Asabuki & Clopath, 2024; Berkes et al., 2011). Sniff fields may provide a reference signal allowing animals to use temporal structure in odor-evoked activity and explain how animals can perceive the timing of sniff-locked optogenetic stimuli (Ackels et al., 2021; Chong & Rinberg, 2018; Hopfield, 1995; Kepecs et al., 2006; Lewis et al., 2021; Li et al., 2014; Powers, 1973; Schaefer & Margrie, 2007; Smear et al., 2011, 2013). Place fields in OB may help unify odor-driven activity with putative internal models instantiated in hippocampus and elsewhere (Buzsáki, 2019; Eichenbaum & Cohen, 2014; Jacobs, 2012; Nieh et al., 2021; O’Keefe & Nadel, 1978; Sheffield & Dombeck, 2015; Sugar & Moser, 2019; Tolman, 1948).

What mechanisms may generate sniff fields and place fields in the olfactory bulb? The olfactory epithelium is mechanically stimulated by airflow, which shapes the activity of olfactory bulb neurons (Buonviso et al., 2006; Díaz-Quesada et al., 2018; Grosmaitre et al., 2007; Iwata et al., 2017). In addition to its feedforward inputs, the olfactory bulb receives centrifugal inputs from neuromodulatory centers and cortical areas (Brunert & Rothermel, 2021; Chen & Padmanabhan, 2022; Kapoor et al., 2016; Linster & Cleland, 2002; Nogi et al., 2020; Reinert & Fukunaga, 2022; Shepherd & Greer, 1998; Soria-Gómez et al., 2014; Sullivan et al., 1989; Zak et al., 2024). Our findings encourage follow up experiments to silence sources of centrifugal innervation of OB to test their impact on sniff and place representations in the bulb.

These results contribute to a body of literature demonstrating that primary sensory areas are modulated by behavior-related information. Spontaneous behaviors, including those unrelated to sensory-guided tasks, describe a significant amount of variation in neural recordings from primary sensory areas (Flossmann & Rochefort, 2021; Long & Zhang, 2021; Mertens et al., 2023; Musall et al., 2019; Parker et al., 2020; Saleem & Busse, 2023; Stringer et al., 2019). The olfactory bulb has been shown to be modulated by reward and task contingencies, inhalation, and other aspects of cognition (Doucette & Restrepo, 2008; Freeman, 1978; Kay et al., 1996; Lindeman et al., 2023; Rojas-Líbano et al., 2014; Zak et al., 2024). As in other systems and species, these representations may be multiplexed by cells to adaptively support sensory coding (Fairhall et al., 2001; Fusi et al., 2016; Panzeri et al., 2010; Weber et al., 2019). Our findings underscore the value of studying sensory systems within more naturalistic behavioral paradigms in which animals are released to perform the repertoire of behaviors in which these sensory systems participate (Buzsáki, 2019; Krakauer et al., 2017; Miller et al., 2022). The importance of active sampling mandates a continued emphasis on detailed observation and quantification of behavioral structure (Bialek, 2022; Marshall et al., 2021; Mazzucato, 2022; Weinreb, Osman, et al., 2024; Weinreb, Pearl, et al., 2024).

## Methods

### Animal housing and care

All procedures were conducted in accordance with the ethical guidelines of the National Institutes of Health and were approved by the Institutional Animal Care and Use Committee at the University of Oregon. Animals were maintained on a reverse 12/12 h light/dark cycle. All recordings were performed during the dark phase of the cycle. Mice were C57Bl6/J background and were 8–12 weeks of age at the time of surgery.

### Surgical procedures

Animals were anesthetized with isoflurane (3% concentration initially, altered during surgery depending on response of the animal to anesthesia). Incision sites were numbed prior to incision with 20 mg/mL lidocaine.

Thermistors were implanted between the nasal bone and inner nasal epithelium (Findley et al., 2021). A custom titanium head bar and Janelia micro drive were implanted.

For olfactory bulb array implantation, we administered atropine (0.03 mg/kg) preoperatively to reduce inflammation and respiratory irregularities. Surgical anesthesia was induced and maintained with isoflurane (1.25–2.0%). Skin overlying the skull between the lambdoid and frontonasal sutures was removed. A rectangular window was cut through the skull overlying the lateral half of the left bulb for insertion of the recording array. The array was lowered to a depth of 1mm and cemented in place with Grip Cement. For hippocampus electrode implantation, an array of 8 tetrodes was inserted vertically through a small, 1mm2 craniotomy overlying the dorsal CA1 cell field of the left hemisphere. To minimize postoperative discomfort, Carpofen (10 mg/kg) was administered 45 minutes prior to the end of surgery. Mice were housed individually after the surgery and allowed 7 days of post-operative recovery.

### Behavioral recordings

Mice were restrained by head fixation then placed in a 15 cm by 40cm behavioral arena. After a period of head-fixation, mice were released to move around the arena, without explicit training or reward structure, while breathing (sampling rate 1kHz), neural data (sampling rate 30kHz), and video (frame rate 100 Hz) were recorded. In a subset of sessions the mice were recorded for 20 min and then the floor was rotated 180 deg and the mice were recorded for an additional 20 min.

The mice were imaged from below to reduce errors due to cable and implant obstruction. A one-direction privacy film was placed on the floor to prevent mice from viewing the open platform which could introduce confounds such as fear responses. A transparent removable flooring was placed directly over this to allow rotation.

We record sniffing using intranasally implanted thermistors (TE Sensor Solutions, #GAG22K7MCD419), amplified initially with custom-built op amp (Texas Instruments, #TLV2460, circuit available upon request) and then a CYGNAS, FLA 01 amplifier fed into the analog input of an open ephys box.

### Pose estimation

The location of the head, center of mass, and base of tail of the mouse were tracked via SLEAP (Pereira et al., 2022). A random set of 1000 frames were hand labeled and compared to the assigned head location to assess error rates. Movement speeds are calculated from the distance the head travels per unit time, smoothed with a 1 s Savitsky-Golay filter. In addition, a head speed limit was placed on the resulting tracking data (10 pix/s = cm/s), and violations were smoothed by linear interpolation.

### Electrophysiology

Following a 3 day recovery period post surgery mice were head fixed and the custom microdrive was advanced to the regions of interest (ROI) while recording. Either Si probes (Diagnostic Biochips P-64-7) or a custom implanted array of 8 tetrodes passed in pairs through 4 linearly-aligned 27-gauge stainless steel hypodermic tubes. Tetrodes were made of 18 µm (25 µm coated) tungsten wire (California Fine Wire). Once the ROI was reached a minimum of 24 hours was allowed prior to data collection to increase recording stability.

Data were acquired via a 128-channel data acquisition system (RHD2000; Intan Technologies) at a 30 kHz sampling frequency and Open Ephys software (http://open-ephys.org). A camera positioned 90cm above the arena floor was used to recording movement around the arena with Bonsai video acquisition software (http://bonsai-rx.org).

Custom Bonsai code was used to align the TTL triggers from the camera frames, the sniff, and the electrophysiology recording captured with no filters applied in the OpenEphys software.

### Spike and sniff data preprocessing and inclusion criteria

Analysis of spikes and sniffing were performed in MATLAB. Electrophysiological data were preprocessed via Kilosort, Phy2, and custom software. Inhalation and exhalation times were extracted by finding peaks and troughs in the temperature signal after downsampling to 1000 samples per s, and smoothing with a 25 ms moving window. All sniffs’ instantaneous frequencies are inverse intersniff intervals. Sniffs with instantaneous frequencies greater than 17 and less than 0.5 sniffs per s were excluded from the analysis. Autocorrelations of sniff frequency and speed and their cross correlation were calculated after mean subtraction and de-trending.

Single units were curated with criteria of 5% refractory period violations (refractory period = 1.5 ms) and an amplitude loss cutoff of 10%. Amplitudes were calculated by first calculating the mean spike waveform on the channel giving the largest spike amplitude and finding its peak and trough times. Then, for each spike time, amplitude was calculated as the difference between the peak and trough times of the mean. The cutoff criterion was this amplitude being less than or equal to zero, so that the fraction of lost spikes can be estimated. This criterion greatly reduces the potential of significant electrode drift over the recording.

### Neuronal population similarity analysis (Matlab)

For population analysis with respect to breathing rhythms (Fig 2), we first calculated each unit’s firing rate time series in 5 s bins, normalized to scale those values between 1 and 0, and smoothed with a 15 s Savitzky-golay filter. For visualization, the units were then sorted according to k-means clustering on the earth movers distances between all units’ time series. These were then colored according to the spike rate colormap used throughout the paper (see below). The resulting matrix is displayed beneath the HMM state-colored sniff frequency plot (Fig 2B). To calculate the similarity matrix over time, we took the cosine distance between all time bins’ population vector (Fig 2B). To compare the neuronal population similarity to the behavioral HMM states, for each session we built a state similarity matrix with the mean cosine distance for comparing every combination of states. To determine whether the distance matrices differed from the prediction of a nonsense correlation null hypothesis, we calculated the state similarity matrix between the neural population vectors when the behavioral HMM states were circularly shifted for the number of 5 s bins in the entire session minus two bins on either side for padding (sessions varied from approximately 30 to 90 minutes). For each session the state similarity matrix values were converted to the number of standard deviations between the real value and the mean of the circular shift null distribution. These similarity matrices were then averaged within animals, and a grand mean was calculated across the within-animal means (Fig 2C). To assess how well the behavioral HMM clustered the neural population vectors, we calculated a silhouette score for the free-moving period of each session. These were then expressed as the number of standard deviations from the mean of a circular shift null distribution.

### Sniff field visualizations (Matlab)

To depict the relationship between individual unit activity and breathing rhythms, we devised sniff fields to portray the relationship between inhalation latency, sniff frequency, and spike probability (Fig 3). For every spike, we assigned a latency as the difference between the spike time and the nearest preceding inhalation time (250 linear spaced bins between 0 and 500 ms), and a frequency as the inverse duration of that sniff (250 log2 spaced bins between 2 and 13 sniffs per s; sniffs with instantaneous frequencies less than 2 were assigned to bin 1, and those greater than 13 to bin 250). Then, to build the joint distribution, we calculated the latency histogram for all spikes that occurred in sniffs within a given frequency bin. We then smoothed the resulting two dimensional matrix with a gaussian filter of width 6. We acknowledge that this smoothing is a questionable choice, given that the latency and frequency axes are in different units, but we nevertheless smoothed in order to reduce the perceptual artifacts of square bins and for aesthetic purposes. These sniff fields were then colored according to the spike rate colormap used throughout the paper (see below).

To analyze sniff fields across the population (Fig 4), we calculated the joint distributions at lower resolution (latency: 30 linear spaced bins between 0 and 300 ms; frequency: 30 log2 spaced bins between 1.75 and 14 sniffs per s). We then took the max projections along the latency and frequency axes for each unit to reduce their information to two “profiles“: latency and frequency. To select for only units with significant selectivity, we then included only those units for which a GLM incorporating the single-variable profiles’ parameter improved the predictions of a latency/frequency model (see below). To compare and cluster these profiles, we calculated the earth movers distance between each pair of profiles (normalized between 0 and 1), and sorted them into types by k-means clustering. We imposed two clusters on the latency profiles and three clusters on the frequency profiles. Including additional clusters did not appreciably change the results. To display these profiles across the population, we stacked the units’ profiles vertically, separated by cluster and sorted by selectivity (the mean of the profile after normalization). These profile stacks were then colored according to the spike rate colormap used throughout the paper (see below).

### Place field visualizations and analysis (Matlab)

To visualize and test the relationship between spiking and place, we assigned each spike to the simultaneous head’s position estimate in a 12 by 5 array of spatial bins (approximately 3 cm squared). Place fields were calculated as the two dimensional distribution of spike positions divided by the occupancy distribution. For visualization, these 12 by 5 maps were scaled to 1200 by 500 pixels and smoothed by a 71 pixel gaussian (about 2 cm squared). This smoothing is intended to reduce the high spatial frequency artifacts resulting from square bins and for aesthetic purposes. These place fields were then colored according to the spike rate colormap used throughout the paper (see below).

To assess the significance of place selectivity, we used the traditional measure of Spatial Information (Skaggs and McNaughton, 1992). The slow variation in position and spike rate time series raise the danger of a nonsense correlation in this metric. Furthermore this metric does not work well for units with high ongoing firing rates. For these reasons, we express spatial information as the Significance of Spatial Information (SSI; Stefanini et al, 2020) calculated as the number of standard deviations from a circular shift null distribution.

### Place Decoding

For place decoding, neural data was binned into 200 ms time windows, and the spatial arena was divided into 60 uniform regions arranged in a 12×5 grid. To account for potential variability over the recording duration, each session was divided into 10 intervals of approximately 4-8 minutes each. Within each interval, a modified 10-fold cross-validation was applied: the interval was subdivided into 10 folds, and the model was trained on 9 folds from each interval while testing on the held-out fold. This cross-validation process was repeated across all intervals, allowing the model to leverage data from the entire session for training and spatial predictions. To classify neural activity by location, we adapted the method from Stefanini et al. (2020) using all identified cells. A Support Vector Machine (SVM) classifier with a linear kernel (implemented via svm.SVC in Python) was used to associate firing patterns with location (Cortes & Vapnik, 1995). Input vectors were non-linearly mapped into a high-dimensional feature space, where a linear decision surface (hyperplane) was constructed. This SVM-based approach enabled pairwise classification across each of the 60 regions in the arena. The classifier employed a majority-vote rule across pairwise outputs to determine the most likely location of the animal, yielding an instantaneous position estimate (Bishop, 2006). The decoded position, set as the center of the selected region, was then used to compute the decoding error between actual and predicted locations.

In a subset of the OB sessions, the arena floor mat was rotated by 180 degrees midway through the session, effectively rotating floor-borne scent marks while leaving distal cues unaffected. For analysis, each session was further partitioned into 10 intervals before the rotation and 10 intervals after, creating 20 intervals in total. To establish baseline decoding accuracy within each half, we applied the original place decoding analysis independently for both pre-rotation and post-rotation periods (Pre-Pre and Post-Post Decoding). To test the generalization of spatial encoding across the rotation, the decoder was trained on 9 folds from each of the 10 pre-rotation intervals and tested on the corresponding 1 fold from the 10 post-rotation intervals (Pre-Post Decoding), with the reverse applied for post-rotation training and pre-rotation testing (Post-Pre Decoding). Additionally, we introduced a fictive 180-degree rotation in the predicted trajectory to evaluate the influence of scent-based versus distal cues. In this fictive rotation analysis (Pre/Post-Rotated and Post/Pre-Rotated Decoding), the model trained on pre-rotation intervals was tested on post-rotation intervals using a fictive 180-degree rotation of the predicted trajectory, and vice versa. This analysis allowed us to evaluate the decoder’s reliance on scent-based versus distal cues.

To assess the significance of decoding error within individual sessions and across all floor rotation conditions, we generated a shuffled baseline by circularly shifting the reversal of the position data by a pseudorandom integer within the middle 80% of the session duration. For each of the 10 folds of this circularly shifted data, we computed 10 median decoding errors. These shuffled errors served as a comparison to the true decoding errors, and statistical significance was assessed using a Wilcoxon rank-sum test.

### Nested Generalized Linear Models (GLMs)

We use Poisson generalized linear models (GLMs) to predict spiking activity of each unit based on sniff parameters and place (Hardcastle et al., 2017). Models were assessed using ten-fold cross-validation. Each session is divided into 50 equal sized bins and ten train test splits are performed on five equally spaced test samples to find representative training and testing sets. Statistical model performance was quantified using a Log-Likelihood Increase (LLHi) metric, which is the change in log-likelihood of held out test data under the fit model compared with a null, mean-rate model. This metric is similar to an F-test, but agnostic to penalization terms. To quantify the non-redundant predictivity of each of these parameters, we calculated a relative predictivity index as the LLHi gained from adding a parameter divided by the total LLHi of the full model. Importantly, because of the redundancy between predictive parameters, these relative predictivity indices do not sum to 1.

### Hidden Markov Models (HMMs)

We fit a gaussian HMM to all sessions from all mice to find behavior states present across mice. The HMM models the observed behavioral data as a gaussian random variable with mean and variance dependent on a time-varying latent state. Behavior observations were formatted as a 5 second moving average of nose speed and a 5 second moving average of the distribution of breathing frequencies. HMM parameters are fit using the Expectation Maximization algorithm. Model selection was performed using Bayesian Information Criterion (BIC), which adds a penalty for increasing the number of model parameters to the likelihood of the data under the model. Lower BIC scores are preferred. BIC scores for 1-10 hidden states are reported as the lowest score of 5 random initializations because EM often finds local maximum in the log posterior. Most likely states were assigned to all sessions from the best performing three-state model using the Viterbi algorithm.

### Colormaps (Matlab)

To make continuous colormaps, we started from a rainbow colormap designed to be perceptually uniform (Kovesi, 2015); colorcet.com). To complement the chromatic variation afforded by this colormap with luminance variation, we multiplied its values with a smooth ramp between 0 and 1. This gives a colormap in which the minimum value is black. Because these figures will be displayed on a white background (i.e., on a website or a piece of paper), we prefer to represent the minimum value as white. To make the minimum value white, we subtracted the colormap from 1. We used the resulting colormap to represent sniffs per s, and by exchanging its red and blue values we made a colormap to represent spikes per s.

To make categorical color maps, we sample colors from paintings or other popular images. Colors were sampled from painters Bridget Riley (Figs 1, 2, 4, 5, and 8; Riley, 1990), Barnett Newman (Fig 7; Newman, 1963), and the credits of the television series Twin Peaks (Fig 4; Lynch & Frost, 1990). We provide a simple Matlab script for sampling colors into colormaps that can be saved and used later for plotting.

## Data and Code Availability

Data and code will be made publicly available at the time of publication.

## Acknowledgements

This work was supported by NIH NINDS R01NS123903, NIH NIDCD R01DC018789, and the Simons Collaboration on the Global Brain (SCGB).

## Author Contributions

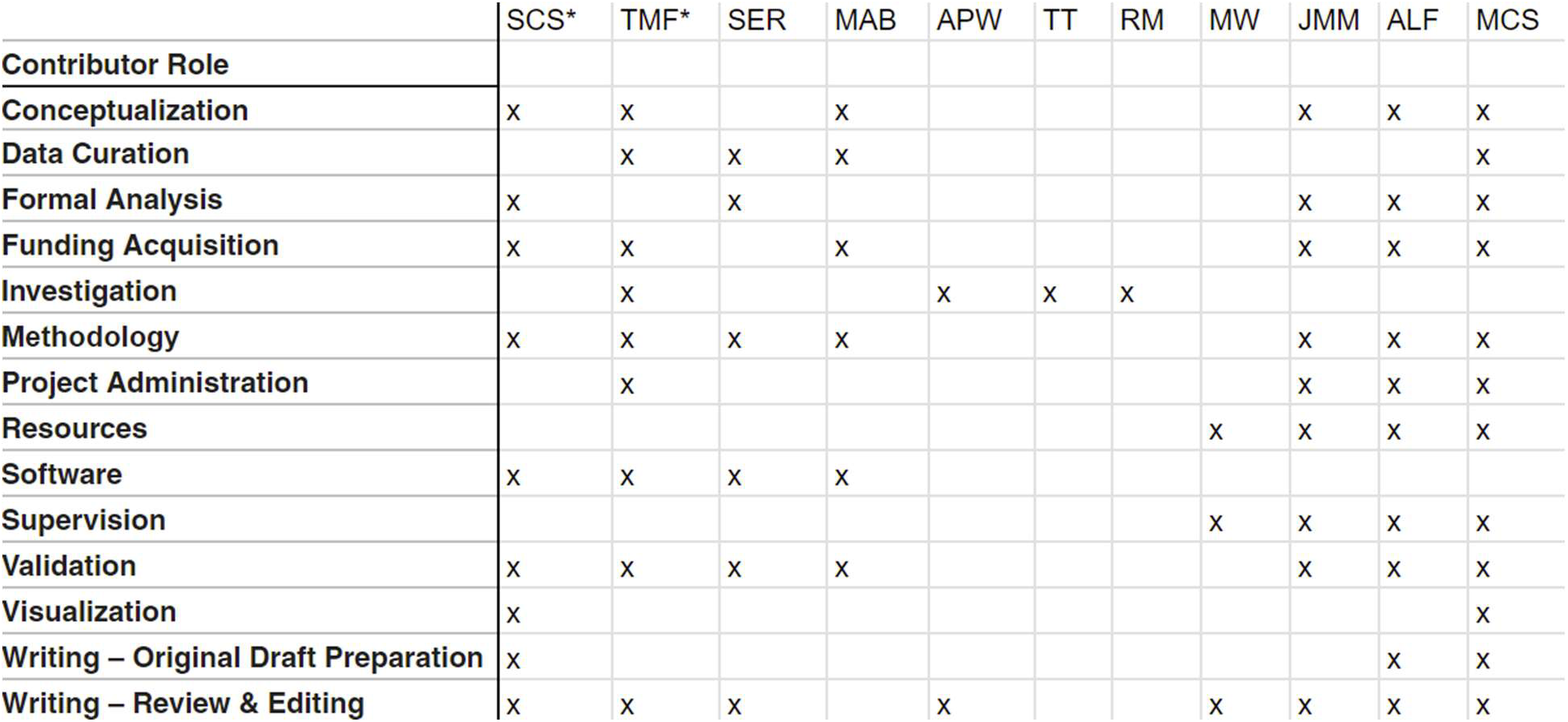

## Supplemental Information

**Figure 1 - Figure Supplement 1:**
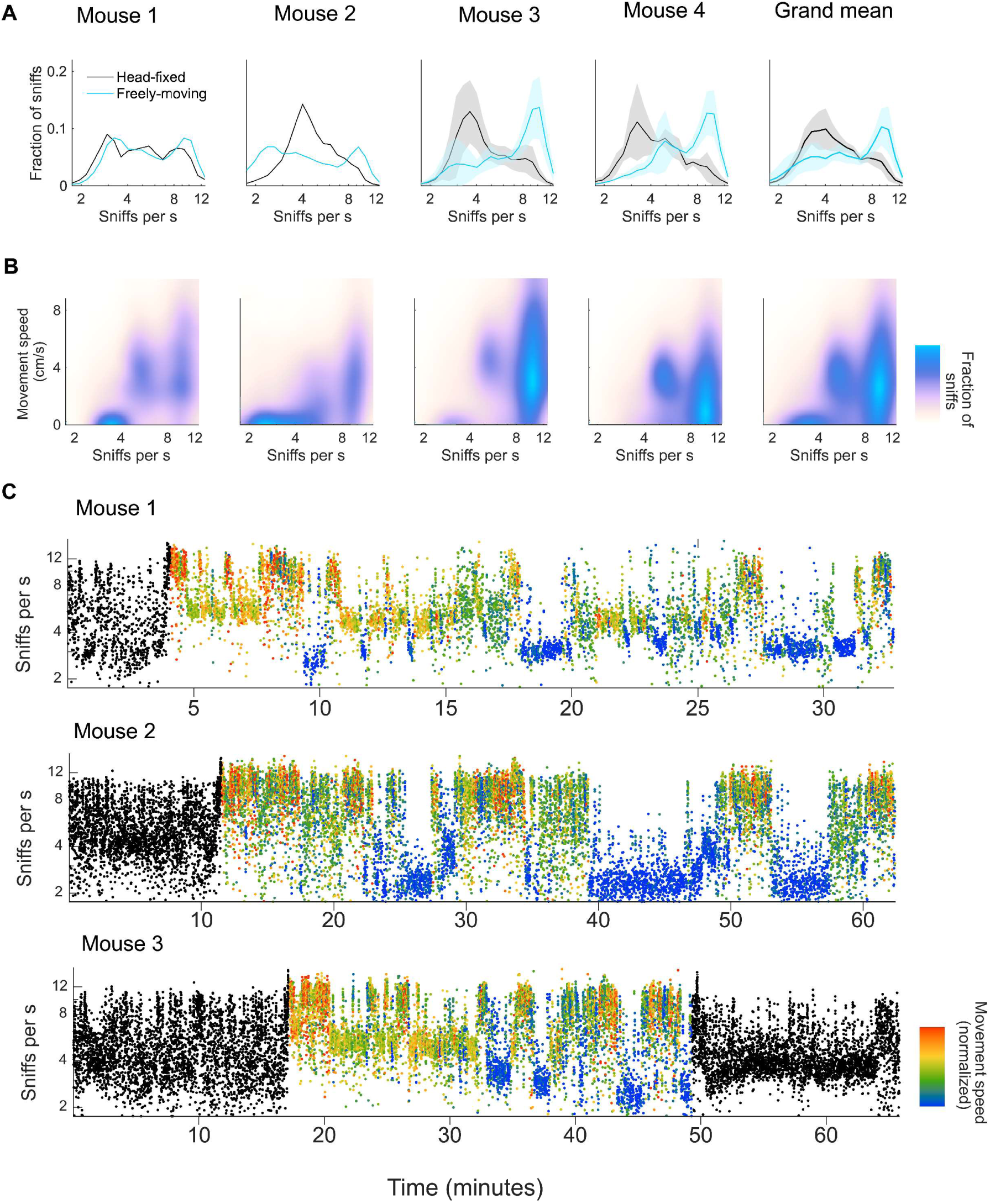
**A.** Histogram of instantaneous sniff frequencies for each individual mouse and grand mean (*n* = 4). Thick lines and shaded regions are mean and ±1 standard deviation. Blue: freely moving; black: head-fixed. **B.** 2D histogram of breathing frequency and movement speed for each individual mouse and grand mean (n=4). **C.** Sniff rasters for three example sessions where each dot indicates an inhalation time with its instantaneous frequency on the vertical axis. Black: head-fixed; Colors based on movement speed during the freely-moving condition.

**Figure 1 - Figure Supplement 2:**
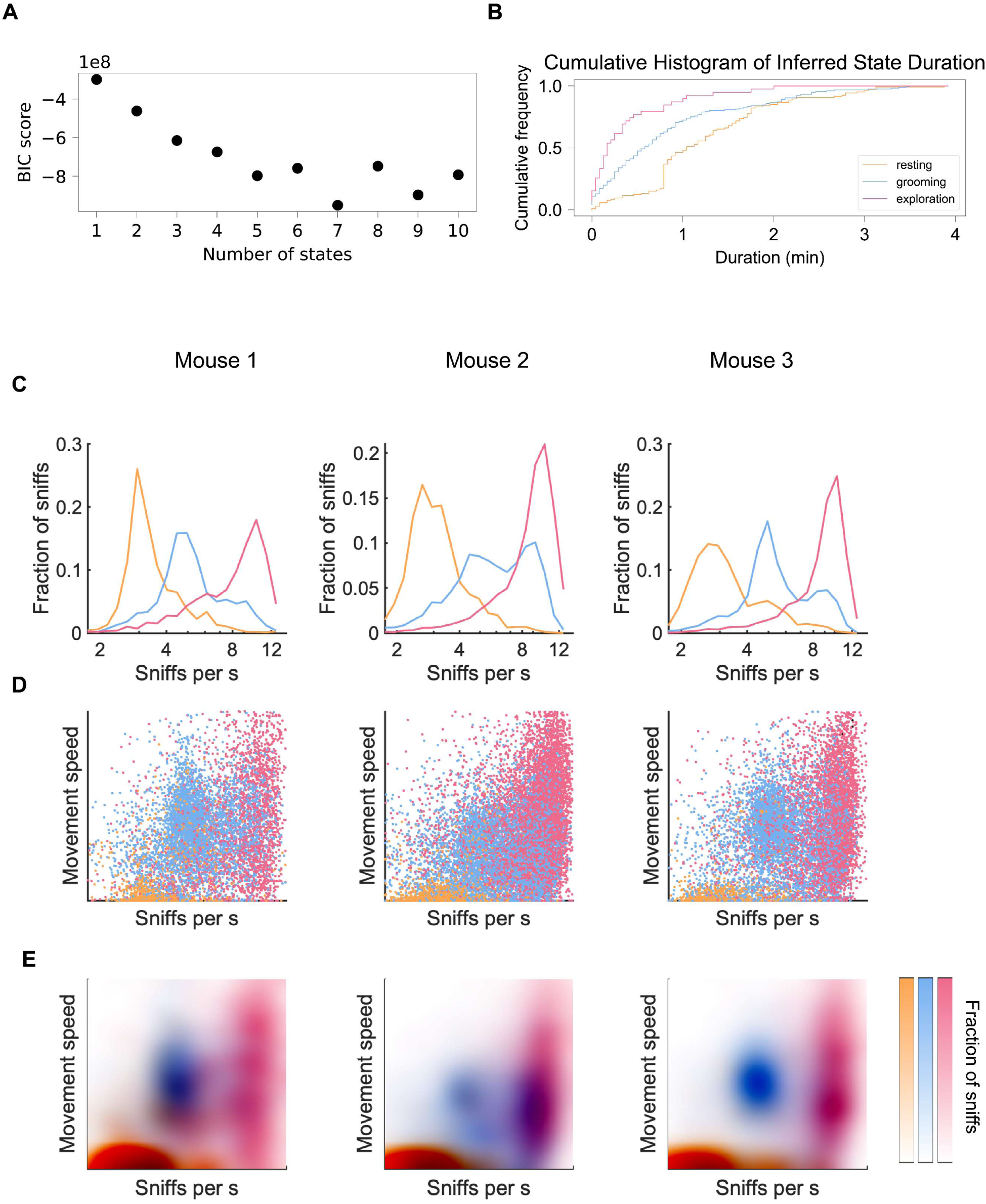
**A.** Bayesian Information Criterion (BIC) scores for HMMs with increasing numbers of hidden states. Scores drop substantially until three states. **B.** Cumulative histogram of inferred state durations across all sessions show that states typically last tens of seconds to minutes. **C.** Sniff Frequency histograms across inferred states for three individual sessions from three mice. **D.** Joint movement speed and sniff frequency scatter plots colored by state assignments for the same three sessions. **E.** Probability density estimates of joint movement speed and sniff frequency for the same three sessions.

**Figure 2 - Figure supplement 1:**
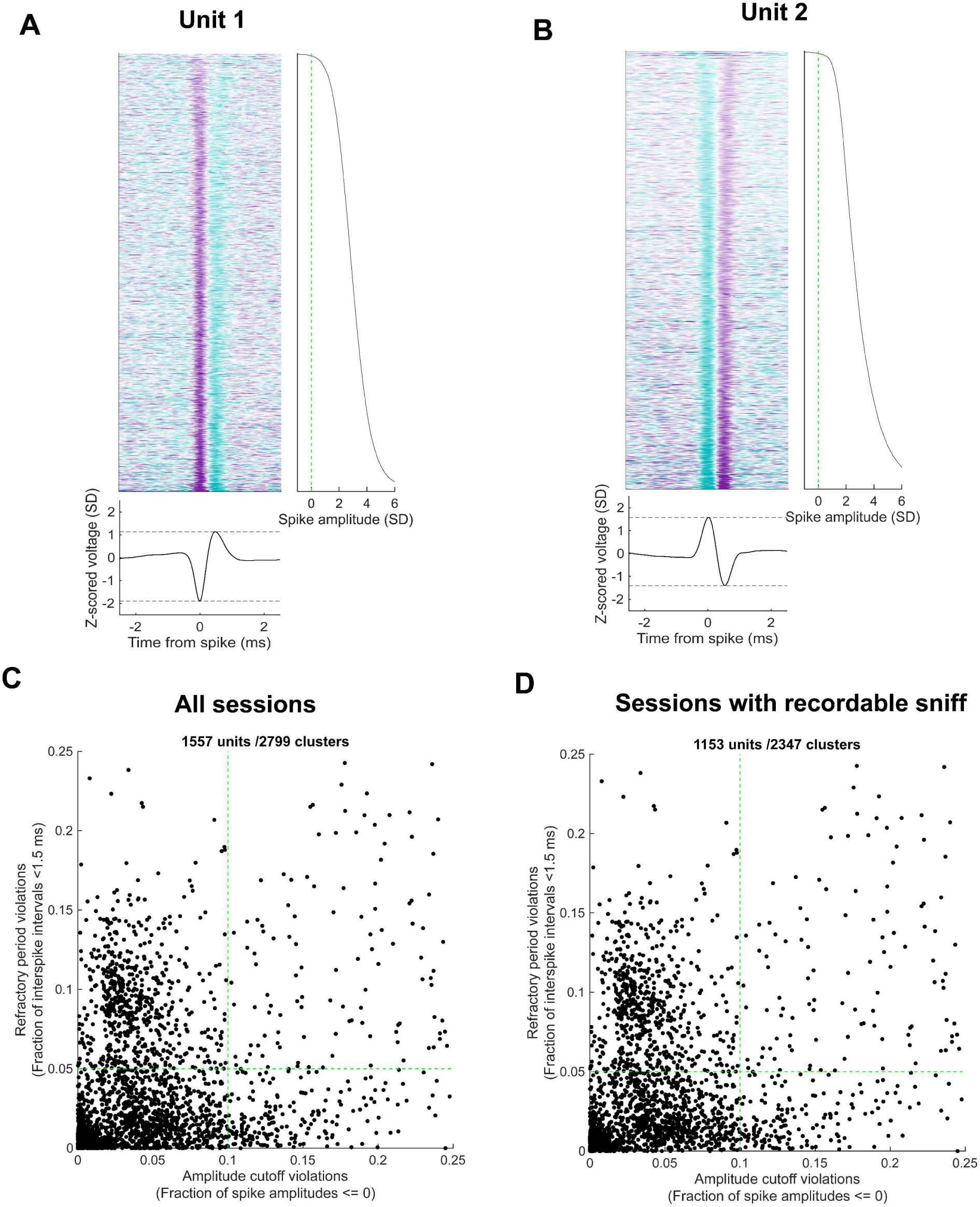
Unit inclusion criteria. **A.** All spike waveforms for an example unit, their corresponding z-scored spike amplitudes, and mean waveform shape. To calculate amplitude on a spike by spike basis, we took the difference between the signal at the peak time and the trough time. **B.** Same as in **A.** for an example unit with positive leading waveform. **C.** Scatter of all cluster’s amplitude cutoff violations and refractory period violations. Green dashed lines show criteria for inclusion (amplitude cutoff violations < 10%, refractory period violations < 5%). **D.** As in **C.** for all clusters in sessions with simultaneous sniff recording.

**Figure 4 - Figure supplement 1:**
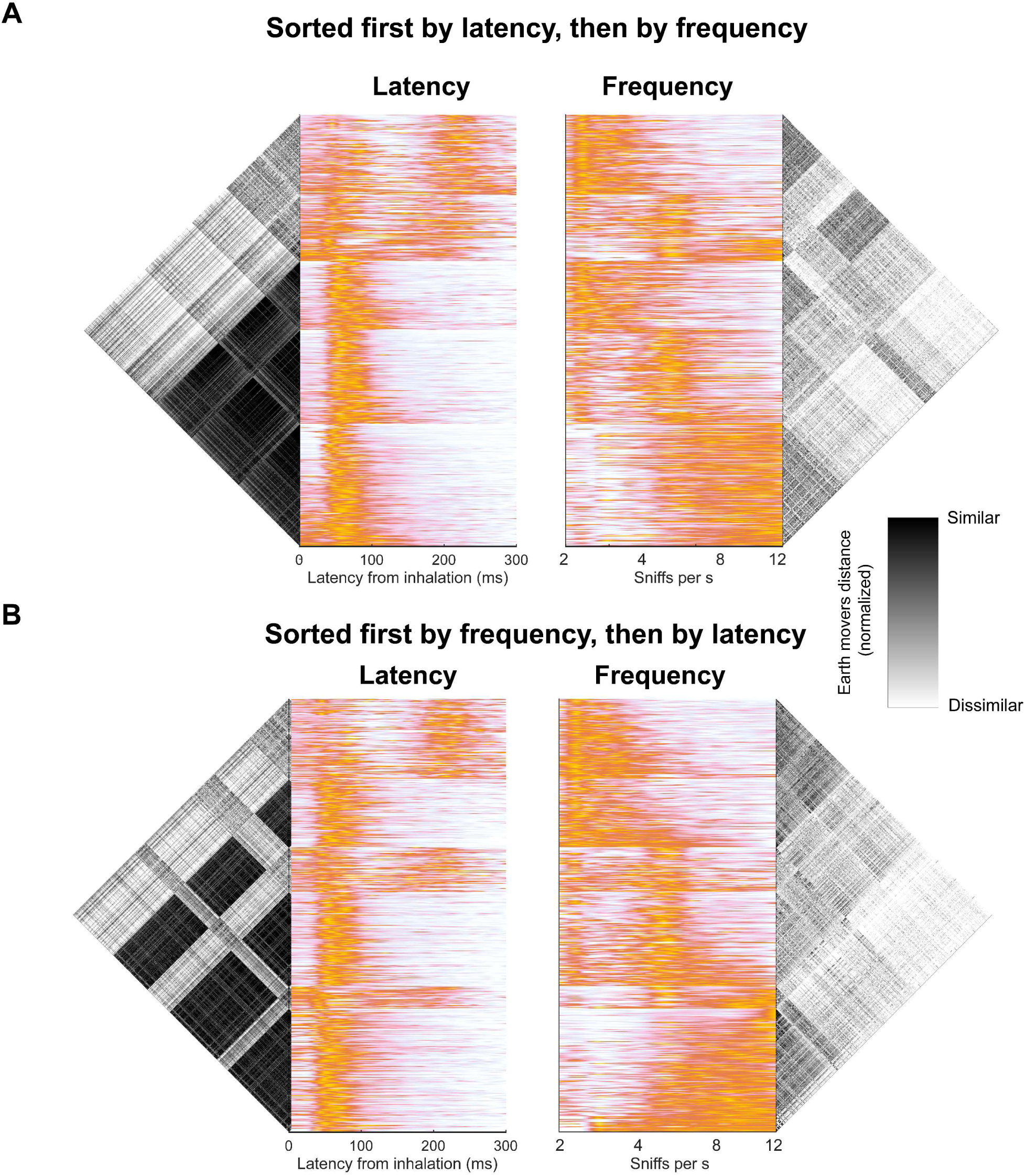
Overlap between sniff latency and frequency clusters. **A.** (Left) SnF latency profiles of all units sorted by latency (Right) SnF frequency profiles separately clustered within latency clusters show a diversity of frequency profiles exist within each functional latency cluster. Earthmovers distance matrices show clustered structure of profiles. **B.** As above, but sorted first by SnF frequency then latency clustered within frequency clusters.

**Figure 8 – Figure supplement 1:**
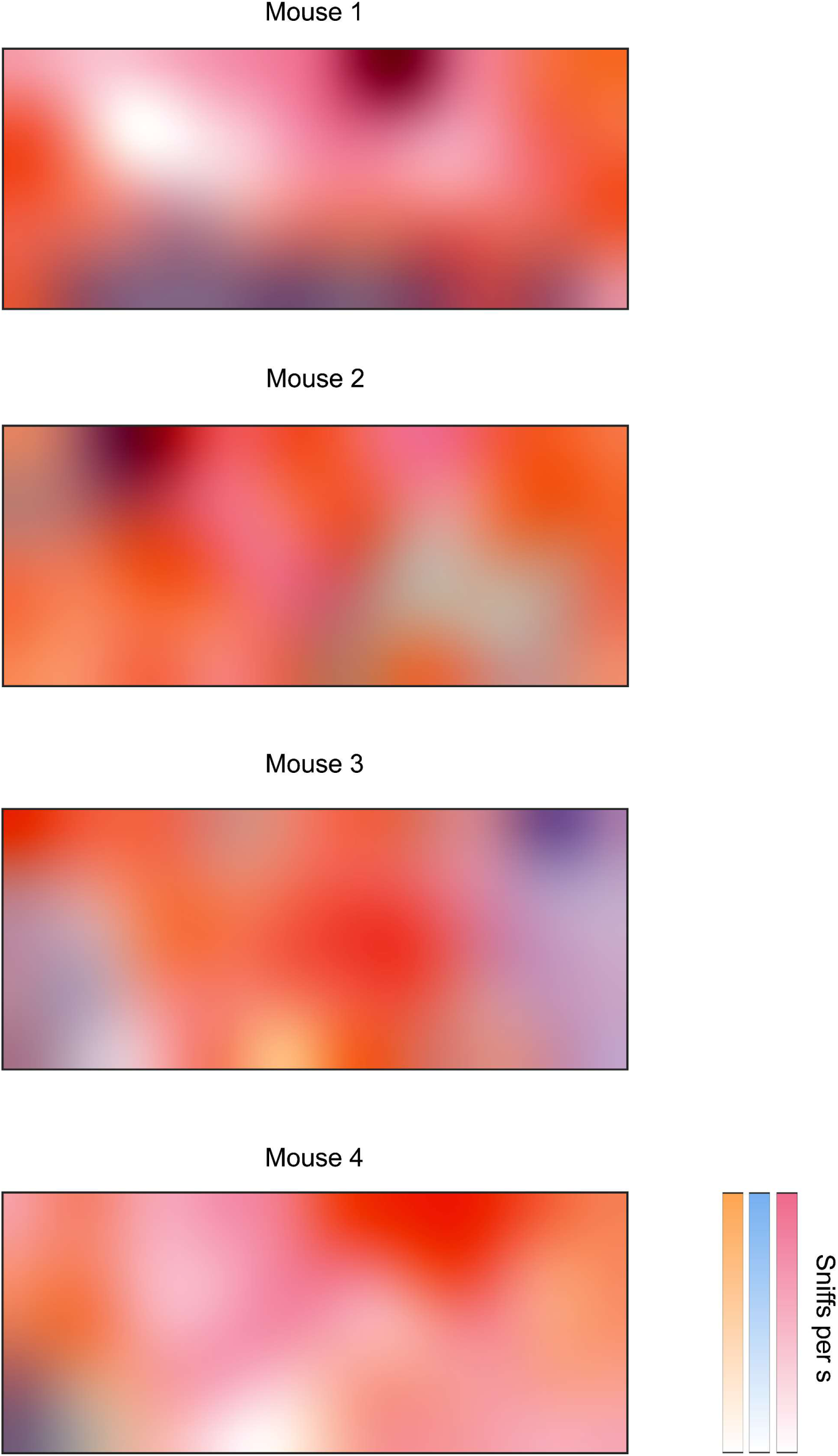
Spatial distribution of behavioral state usage. The spatial distribution of behavioral state usage for each OB mouse. Colormap overlays the usage of each state, normalized by sniffs, into a composite color as in Figure 1 (Figure 1 – Supplemental video 2).

**Figure 1 - Supplemental Video 1: Visualizing and sonifying neurodata (ViSoND) from a representative recording session.** To provide an observable demonstration of neural recordings during freely moving behavior, we developed a tool called ViSoND, in which sniff and spike events are rendered to MIDI notes, so that different events can be identified by different sounds. The top panel shows video of the mouse, the center panel shows the synchronous raw thermistor signal, colored according to the current behavioral HMM state, and the bottom panel animates a population raster that is also synchronous with the behavior video. Inhalation times are indicated visually by peaks in the thermistor signal and sonically by occurrences of a kick drum sample. Visually, spikes from each unit are indicated by marks on each row of the raster plot. Sonically, spikes from each unit are mapped to a different note of a virtual piano. Accessible at https://www.dropbox.com/scl/fi/2kae0qnftw052e9gcn7ig/Sniff_OB_ViSoND.mp4?rlkey=cq926o6ntazlw4oq4dzguwju0&dl=0

**Figure 1 - Supplemental Video 2: 3D color map visualization for distribution-weighted color mixing.** In order to indicate the distributions for three populations, defined by our behavioral states, we developed distribution-weighted color mixing. Here, each state is identified by a basis color from Riley (1990), and multiplied with the distribution of sniffs for a given set of parameters. Colors range from white to full color for a given state, and overlap is indicated by darkening. This color scheme can be conceptualized as a cube, with the three axes defined by the three states (“Explore”, “Groom”, and “Rest”) and represented by their respective basis colors. Video animates sections from this cube along each of the three axes, where each subplot indicates sections taken from a different angle. Accessible at https://www.dropbox.com/scl/fi/qofn0fh593xjy2myqspwa/Figure-1-Extended-video-2.mp4?rlkey=n3w9g7q0w18h81ftxgb64g7br&dl=0

